# Hepatic loss of AATF attenuates MASH by suppressing AKT–mTORC1 signaling and reprogramming lipid metabolism

**DOI:** 10.1101/2024.05.25.595878

**Authors:** Akshatha N. Srinivas, Diwakar Suresh, Prajna Anirvan, Manju Moorthy, Gopalkrishna Ramaswamy, Suma M. Nataraj, Prasanna Kumar Santhekadur, Deepak Suvarna, Shivaram P. Singh, Divya P. Kumar

## Abstract

**Background & aims:** Metabolic dysfunction-associated steatohepatitis (MASH) is a multifactorial disease driven by complex molecular mechanisms. Identifying key regulators is critical for developing targeted therapies. Here, we demonstrate the impact of the loss of apoptosis antagonizing transcription factor (AATF) on hepatic lipid metabolism and MASH progression.

**Methods:** A preclinical mouse model recapitulating human MASH was established by feeding C57Bl/6 mice either a chow diet (CD) or a western diet with sugar water (WD). Hepatic AATF silencing was achieved by tail vein injection of siAATF delivered by adeno-associated virus 8 (AAV8) with the liver-specific thyroxine-binding globulin (TBG) promoter. In addition to histological, biochemical, and molecular biology evaluations, the mechanistic insights were derived from whole transcriptomics and untargeted metabolomics analyses.

**Results:** AAV8-mediated specific knockdown of AATF in hepatocytes significantly reduced body weight, liver weight, and insulin resistance in mice fed with western diet (WD). However, no such effects were observed in mice fed with a chow diet (CD). Further analyses showed reduced liver injury, steatosis, and steatohepatitis in WDsiAATF mice. Transcriptomic analysis demonstrated that loss of AATF alleviated cellular stress, inflammation, and fibrosis in WD-fed mice. Moreover, AATF silencing led to alterations in lipid metabolism, notably by decreasing hepatic lipogenesis in the WD mice. Interestingly, untargeted metabolomics revealed an increase in the biosynthesis of glycerophospholipids and beta-oxidation of fatty acids in WDsiAATF mice.

**Conclusion:** Our findings reveal a previously unrecognized role of AATF as a central regulator of hepatic lipid metabolism in MASH, acting through the AKT–mTORC1 signaling pathway, and establish its inhibition as a promising therapeutic strategy for metabolic liver disease.

## Background

Metabolic dysfunction-associated steatotic liver disease (MASLD), formerly termed non-alcoholic fatty liver disease (NAFLD), stands as the prevailing chronic liver ailment impacting more than 30% of the adult population worldwide (1,2). It notably afflicts individuals with obesity and type 2 diabetes, emerging as a primary contributor to advanced liver disease and hepatocellular carcinoma (HCC) (3,4). MASLD encompasses a spectrum of advancing steatotic liver conditions, spanning from benign metabolic dysfunction-associated steatotic liver (MASL) to steatohepatitis (MASH), presenting with varying degrees of liver fibrosis that could advance to cirrhosis (5). MASLD not only heightens the likelihood of liver-related complications such as cirrhosis, end-stage liver disease, and hepatocellular carcinoma (HCC) but also escalates the risk of multiple extraneous manifestations, including cardiovascular disease (CVD), chronic kidney disease (CKD), and specific forms of extrahepatic cancers (6). As MASLD progresses to the end stage of liver disease, treatment options become increasingly limited, often leading to consideration for liver transplantation (7). Early detection and intervention are, therefore, paramount. While lifestyle modifications remain fundamental to managing the condition, pharmacological treatments can complement these interventions (8,9). Despite numerous clinical studies, the pathogenesis of MASH remains elusive, and the mechanisms underlying its occurrence and progression are not yet fully understood (10). Therefore, elucidating the molecular pathways involved in MASH pathogenesis is crucial for identifying potential therapeutic targets.

MASH manifests with the accumulation of lipids in the liver, stemming from disturbances in both lipid uptake (from circulating sources or de novo synthesis) and lipid disposal (via fatty acid oxidation or secretion of triglyceride-rich lipoproteins) (11). This disruption leads to the accumulation of lipotoxic compounds, cellular stress, injury, and fibrosis (11,12). The regulation of lipid metabolism is complex and multifaceted, involving various factors like membrane transporters, metabolic enzymes, and transcriptional regulators (13). Glycerophospholipids, a subclass of phospholipids that are the major constituents of cell membranes, play a crucial role in lipid metabolism. Alterations in the composition of glycerophospholipids have been reported in individuals with MASH (14,15). Phosphatidylcholine (PC), phosphatidylethanolamine (PE), phosphatidylserine (PS), phosphatidylinositol (PI), and cardiolipin (CL) are a diverse array of glycerophospholipids that are well-studied for their roles in cell signaling and lipid metabolism (15). Interestingly, glycerophospholipids are involved in inflammatory, oxidative stress, and bile acid metabolism pathways, which are key drivers of disease pathogenesis in MASH (16). Notably, glycerophospholipids are crucial for maintaining the integrity of the mitochondrial membrane, which is essential for various mitochondrial functions, including the beta-oxidation of fatty acids (17). It is observed that, in MASH subjects, diminished β-oxidation of fatty acids in the liver fails to adequately mitigate the lipotoxic effects of harmful lipids (18). Disruption of β-oxidation pathways leads to lipid accumulation in the liver, increased oxidative stress, and inflammation, all of which contribute to the progression of the disease (19). Thus, glycerophospholipid levels and efficient β-oxidation of fatty acids in the liver contribute to the protection and alleviation of MASLD.

Apoptosis antagonizing transcription factor (AATF) is a multifaceted transcription factor recognized for its pivotal involvement in essential cellular functions (20). It serves as a dynamic regulator of fundamental cellular processes, including transcriptional control, regulation of the cell cycle and apoptosis, ribosome biogenesis, and response to cellular stress (20). AATF protects cells from oxidative stress, ER stress, DNA damage, and hypoxia by modulating the cell cycle, autophagy, and apoptosis (21). The widely studied role of AATF is the regulation of apoptosis, exerted by the activation of antiapoptotic pathways and the inhibition of proapoptotic pathways (22). Pathologically, AATF has been implicated in numerous cancer types, including leukemia, breast cancer, neuroblastoma, lung cancer, and colon cancer (23–25). Our previous investigations have revealed elevated levels of hepatic AATF in HCC, where it plays a significant role in promoting tumor angiogenesis (26). Studies have also demonstrated the inhibitory effects of curcumin on MASH-HCC by downregulating AATF via the KLF4-SP1 pathway (27). We recently investigated the role of the SIRT1-TIMP3-TACE axis in AATF upregulation, with subsequent TACE inhibition resulting in AATF downregulation and the attenuation of MASH-associated liver tumorigenesis (28). However, the mechanistic role of AATF in the pathophysiology of MASH remains poorly understood.

Despite growing evidence implicating metabolic and inflammatory pathways in MASH, the transcriptional regulators that coordinate these complex responses remain poorly defined. Here, we demonstrate for the first time that hepatocyte-specific loss of AATF ameliorates MASH through metabolic reprogramming, including enhanced β-oxidation, suppressed lipogenesis, and restoration of glycerophospholipid homeostasis. By integrating transcriptomic and untargeted metabolomic profiling with in vivo AAV8-mediated gene silencing, our study provides a novel mechanistic framework linking AATF to MASH pathogenesis and offers a previously unexplored therapeutic target for this widespread liver disease.

## Methods

### Animal study

Male C57Bl/6NCrl mice aged 6 weeks procured from Hylasco Biotechnology Pvt. Ltd. (Charles River Laboratories, Hyderabad, Telangana, India) were used for the study. All the mice were accommodated in the central animal housing facility at the Center for Experimental Pharmacology and Toxicology, JSS AHER, following a 12-hour light-dark cycle, consistent with previously established protocols (59). Following one week of acclimatization, study protocols were commenced. All animal-related procedures adhered to the guidelines sanctioned by the Committee for the Purpose of Control and Supervision of Experiments on Animals (CPCSEA), Govt of India (JSSAHER/CPT/IAEC/118/2022).

### Diet-induced MASH model

After a week of acclimatization, the mice were fed with either a chow diet normal water (CD) or a western diet (21% fat, 41% sucrose, and 1.25% cholesterol by weight, Research Diet, Inc. USA) along with sugar water (WD) containing 18.9 g/L d-glucose and 23.1 g/L d-fructose (HiMedia Laboratories) ad libitum for 24 weeks. Mice were continuously monitored for food intake, water intake, and body weight. After 24 weeks, mice were euthanized, and blood and tissues were collected for further histological, biochemical, and molecular analysis.

### AAV expression system

Liver gene expression was modulated using AAV-mediated gene transfer technology, as previously defined (60). The siRNA sequences targeting AATF were incorporated into an adeno-associated virus (AAV) vector under the regulation of a liver-specific thyroxine-binding globulin (TBG) promoter. AAV8-TBG-siAATF and AAV8-TBG-siControl constructs were generated by packaging into serotype 8, titrated by Applied Biological Materials Inc. (ABM). The siRNA construct design mimics the behavior of shRNA due to its vector-based expression system. The siRNA sequence is cloned into a plasmid vector under the control of a dual convergent promoter system, U6 and H1, which drives the transcription of both strands of DNA to form a shRNA-like structure within the cell. This technology enhances the stability of siRNA, enabling prolonged gene silencing. Henceforth, the one-time insertion of siRNA in the study provided a stable and sustained expression.

### Hepatocyte-specific silencing of AATF in experimental MASH

The mice were fed either CD or WD. After 12 weeks of dietary intervention, mice were injected via the tail vein with 1×10^11^ genome copies of AAV8 serotype particles and randomized into four groups: (i) CD with AAV8-TBG-siControl, (ii) CD with AAV8-TBG-siAATF, (iii) WD with AAV8-TBG-siControl, and (iv) WD with AAV8-TBG-siAATF. The same dietary regimen was followed until the end of 24 weeks. Throughout the study, mice were continuously monitored for food intake, water intake, and body weight. The GTT and ITT were performed during the study period. After 24 weeks, mice were euthanized under fasting conditions, followed by blood and tissue collection for further analysis.

### Isolation of primary mouse hepatocytes

Primary hepatocytes were isolated from the livers of mice injected with AAV8-TBG-Null and AAV8-TBG-siAATF following previously established protocols (60). Briefly, mice were anesthetized, and the inferior vena cava was cannulated. Liver perfusion was carried out using pre-warmed perfusion buffer, followed by digestion buffer containing 25 µg/ml Liberase (Sigma-Aldrich). The isolated hepatocytes were then filtered through a 70-micron cell strainer (HiMedia) and purified via centrifugation in DMEM low-glucose medium (HiMedia Laboratories) at 50g for 5 minutes at room temperature. Percoll (Sigma Aldrich) gradient separation was employed to isolate hepatocytes from nonparenchymal cells. The isolated cells were suspended in DMEM supplemented with 10% FBS and 100 U penicillin-streptomycin. Collagen-coated plates were utilized for cell culture in William’s E medium (HiMedia Laboratories).

### Glucose and Insulin Tolerance Tests

Mice were fasted overnight, or 5-6 hours for GTT and ITT, respectively. The basal level of blood glucose was measured using a glucometer (Roche). Furthermore, mice were administered glucose orally (1 g/kg body weight) and insulin intraperitoneally (0.75 U/kg body weight). Venous tail blood samples were taken at various intervals between 0 and 120 minutes to measure blood glucose levels.

### Biochemical analysis

Blood collected from the fasting mice was centrifuged at 1500 g for 15 min at 4°C. The serum levels of AST, ALT, total cholesterol (TC), total triglycerides (TG), and LDL-cholesterol were measured using commercial kits (Monlab tests) according to the manufacturer’s protocol.

### Histological analysis

Liver tissues were subjected to formalin (Sigma Aldrich) fixation, and paraffin blocks were prepared. 4µm tissue sections were used for hematoxylin and eosin staining to assess the liver morphology. For pathological grading, stained liver sections were scored by pathologists according to the MASLD activity score, which comprises the sum of steatosis (0-3), lobular inflammation (0-3), and hepatocyte ballooning (0-2). A cumulative activity score of 5 or greater was indicative of MASH (61,62).

### Oil Red O staining

Oil Red O staining was performed using frozen liver tissues. Cryosections, 12 μm thick, were fixed with 10% formaldehyde (Sigma Aldrich) for 5 minutes and then washed with distilled water. Further, the slides were incubated with 60% isopropanol (Sisco Research Laboratories) and Oil Red O solution (Sigma Aldrich) for one hour. After rinsing off excess stain with distilled water, nuclei were counterstained with hematoxylin (HiMedia Laboratories). The slides were then mounted, and images were captured using the Leica DM570 microscope (Leica Microsystems). For quantitative analysis, the areas of lipid droplets were quantified using ImageJ software.

### Picrosirius red staining

The formalin-fixed paraffin-embedded (FFPE) liver tissue sections were subjected to deparaffinization and rehydration using xylene (HiMedia), followed by decreasing percentages of ethanol. The slides were first stained with hematoxylin for 2 minutes, followed by incubation with the picrosirius red solution (Sigma Aldrich) for 1 hour. Slides were washed in 0.5% acidified water and mounted. The images were captured using the Leica DM570 microscope (Leica Microsystems) and quantified using ImageJ software.

### ELISA

Serum insulin and TNFα levels were quantified using mouse ELISA kits (Krishgen Biosystems) according to the manufacturer’s instructions.

### Immunohistochemistry

Immunohistochemistry was performed according to standard protocol. Briefly, after deparaffinization and rehydration, FFPE liver tissues were subjected to antigen retrieval with citrate buffer (pH 6) at 94°C for 15 minutes. Following this, the tissue sections were blocked with 3% hydrogen peroxide (PathnSitu Biotechnologies) and normal goat serum (Abcam) for 2 hours. Primary antibodies (AATF; Sigma Aldrich, 1:500, F4/80; Cell Signaling Technologies, 1:100, and Desmin; Cell Signaling Technologies, 1:500) were used to stain the respective tissue sections overnight at 4°C in a humidified chamber. The detection of signals was accomplished using the Polyexcel HRP/DAB detection system (PathnSitu, Biotechnologies). Images were captured using the Leica DM570 microscope (Leica Microsystems), and quantitative analysis was conducted using ImageJ software.

### Immunoblot analysis

Liver tissues and cell pellets were dissolved in RIPA buffer (Sigma Aldrich) containing protease/phosphatase inhibitor cocktail (Thermo Fischer Scientific). The lysates were obtained by centrifugation at 12000 rpm at 4^0^C for 20 minutes. Protein concentration was estimated by Bradford’s (Bio-Rad) protein estimation method. Equivalent amounts of protein were resolved in SDS-PAGE, followed by transfer into the nitrocellulose membrane (Bio-Rad). Blots were blocked in 5% nonfat dry milk for 1 hour and probed with primary antibodies of interest (anti-AATF-Sigma Aldrich 1:1000, anti-pAKT, anti-AKT, anti-pmTOR, anti-mTOR, anti-Raptor, and anti-β-actin - Cell Signaling Technologies 1:1000, anti-pJNK, anti-JNK, anti-pERK1/2, and anti-ERK1/2 – Santa Cruz Biotechnologies 1:100) overnight at 4°C and secondary antibodies (Cell Signaling Technologies) for 2 hours at room temperature. Blots were visualized using Western Bright ECL HRP substrate (Advansta) in the UVitec

Alliance Q9 chemiluminescence imaging system and analyzed for densitometric measurements in ImageJ software. The band intensity was normalized with the respective endogenous control.

### RNA isolation, cDNA synthesis, and Quantitative real-time PCR

Total RNA was extracted from frozen live tissues or cell pellets using TRIzol reagent (Sigma-Aldrich) according to the manufacturer’s instructions. RNA was quantified using a Nanodrop spectrophotometer and reverse-transcribed using the Verso cDNA synthesis kit, following the manufacturer’s protocol (Thermo Fisher Scientific). Relative gene expressions were assessed using quantitative real-time PCR (qRT-PCR) using the Rotor-Gene Q 5plex HRM system. qRT-PCR was performed in triplicate per gene using the Dynamo Colorflash SYBR Green kit (Thermo Fisher Scientific) with 0.5 μM primers (IDT), and 50 ng of cDNA in a 20 μL reaction volume was used to set up the reaction. The mRNA levels of selected genes were calculated after normalization to β-actin by using the 2^(-ΔΔC(T))^ method. Primer sequences are provided in Table S1.

### RNA sequencing and transcriptomics analysis

RNA extraction was performed using the All-prep DNA/RNA kit (Qiagen). RNA was quantified using a Qubit fluorometer, and RIN was determined with a TapeStation 4150 using HS RNA screen tape. The library preparation was carried out using the SMART-Seq Library Prep Kit. The sequencing was performed using the Illumina NovaSeq 6000 platform with a read length of 150 bp × 2. The reads were mapped to STAR-indexed Mus musculus (GRCm39: GCF_000001635.27) using the STAR aligner version 2.7.9a. The alignment file (sorted BAM) from individual samples was quantified using Feature Counts version 2.0.1, based on the filtered GTF file, to obtain gene counts. These gene counts for Mus musculus were used as inputs to DESeq2 for estimating differential expression.

### Differential gene expression analysis

Differential expression analysis was performed using the Bioconductor package DESeq2 (R, version [latest version]). Genes with read counts of less than 10 across the samples were removed before proceeding with downstream analysis. Raw read counts were normalized by DESeq2. The magnitude (log 2 transformed fold change) and significance (P-value) of differential expression between groups were calculated, and genes with P-values <0.05 were considered significant. We obtained DEGs with the following criteria: P value <0.05 and |log 2-fold change|≥2. PLS-DA analysis was carried out in R using the plsda function in the “mixOmics” package.

### Functional Gene Set Enrichment Analysis

Functional gene set enrichment analysis (GSEA) was performed using the R package ClusterProfiler for Gene Ontology Biological Process (GO BP), Molecular Function (GO MF), and Kyoto Encyclopedia of Genes and Genomes (KEGG) pathways. For multiple test corrections, P-values were adjusted by the Benjamini-Hochberg false discovery rate (BH-FDR). Gene sets with a p-value < 0.05 were considered significantly enriched. Gene sets were downloaded from the MSigDB database (https://www.gsea-msigdb.org/gsea/msigdb).

### Gene signature heatmaps

Normalized expression of leading genes in the enriched cellular stress, inflammation, and fibrosis pathways identified by GSEA was used to generate heatmaps using the pheatmap R package. Glycerophospholipid metabolism, lipid metabolism, and fatty acid degradation pathway gene lists were obtained from the KEGG database.

### AATF PPI network

The Search Tool for the Retrieval of Interacting Genes (STRING version 12.0) (http://string-db.org) was used to systematically screen the top 100 mouse protein interactors of AATF, set with a high confidence threshold of > 0.7. Interactors of AATF were also retrieved from the BioGRID database. Furthermore, the protein-protein interaction network was created using Cytoscape software, where the color of the node represents the log2 fold change of the gene within the comparison of WDsiAATF vs WDC. Interactions between AATF and genes involved in biological processes related to lipid metabolism, as defined by the Gene Ontology, were screened separately within the STRING database. The gene ontology biological process and molecular functions, as well as KEGG pathways enriched by the interactors of the AATF gene, were analyzed and visualized using the ClueGO app within Cytoscape. For all the data related to transcriptomics, P<0.05 was considered to indicate a statistically significant difference. All statistical analysis and visualizations were performed in R v.3.2 unless indicated otherwise.

### Untargeted metabolomics-Metabolite extraction and LC-MS analysis

Metabolites were extracted following previously established protocols (63). Briefly, liver tissues were lyophilized in a vacuum freeze dryer for 24 hours, and 50 mg of samples were homogenized with 50 mM ammonium bicarbonate (3:1 w/v). Samples were extracted in cold methanol by centrifugation at 18,000× g for 15 minutes at 4°C. Supernatants are dried and reconstituted in a 3:1 (v/v) mixture of MeOH and water for LC/MS analysis. The hepatic metabolome was analyzed using a Vanquish UHPLC system (Thermo Fisher) coupled with an Orbitrap Eclipse Tribrid mass spectrometer (Thermo Fisher). Samples were separated on an RP column, 2.1 x 150 mm, 1.8 μm particle size, and 0.3 mL/min flow rate with two eluents for positive and negative polarity modes.

### Metabolomics data analysis

The data processing was done using the Compound Discoverer 3.3 software (Thermo Fisher Scientific). A pre-existing workflow template titled ‘Untargeted Metabolomics with Statistics Detect Unknowns with ID using Online Databases and mzLogic’ was used to identify the differences between samples. For identification, the mass tolerance was 5 ppm, and the retention time tolerance was 0.2 min. The minimum peak intensity was 10,000, and the chromatographic S/N threshold was 1.5. The solvent blanks were used to create exclusion lists, identify background components, and filter these components from the results table. A pooled quality control (QC) sample was used for compound identification only.

To visualize the similarity between the metabolomic profiles of the samples from different groups, principal component analysis (PCA) and heatmap plots were generated. All the studies were conducted based on the compound’s peak areas. Differential metabolite analysis was performed using a student’s t-test, considering p value<0.05 as significant and log2 foldchange ≥ |1| as either an upregulated or downregulated metabolite. The log2 fold change per metabolite was calculated as the ratio between group areas, transformed to a log2 scale, with the group area being calculated as the median of the compound area per sample within the group. The up-or down-regulated metabolites were visualized using volcano plots. P-values were adjusted for multiple testing using the Benjamini-Hochberg correction for false discovery rate.

The GSEA algorithm was applied to the KEGG Mus musculus (House Mouse) pathway library, specifically the lipid superclass and its subclasses. Significant pathways (p < 0.05) identified by the GSEA method were further analyzed. Pathway enrichment against the Small Molecule Pathway Database (SMPDB) was performed separately for upregulated and downregulated metabolites using MetaboAnalyst 6.0. For overlaying the regulation of metabolites involved in the glycerophospholipid metabolism pathway, the compound list and pathway were accessed from the KEGG database and visualized using the Pathview R package. To visualize the metabolic network, MetScape (http://metscape.ncibi.org/) was used, and the Cytoscape plugin was employed to identify the interactions between the metabolites involved in the glycerophospholipid metabolism pathway.

### Quantification and statistical analysis

Statistical analyses were performed using GraphPad Prism 6.0 software (GraphPad, Inc., La Jolla, CA, USA). Data were represented as the mean ± standard error of the mean. Statistical significance was analyzed using Student’s t-test for two groups or analysis of variance (ANOVA) with post hoc Bonferroni correction for multiple comparisons, and p values < 0.05 (*/^#^) or < 0.001 (**/^##^) were considered significant.

## Results

### Loss of hepatic AATF ameliorates western diet-induced insulin resistance, liver injury, steatosis, and steatohepatitis

To assess the role of AATF in MASH, we targeted AATF expression in mouse hepatocytes using a liver-directed adeno-associated virus serotype 8 (AAV8) vector driven by the thyroxine-binding globulin (TBG) promoter. Mice were fed a chow diet with normal water (CD) or a western diet with sugar water (WD). After 12 weeks, they were administered either AAV8-TBG-siControl or AAV8-TBG-siAATF (The stable gene silencing effect is consistent with the engineered siRNA construct’s shRNA-like behavior, driven by a dual-promoter expression system-*Detailed in the Methods section*) via tail vein injection and monitored for an additional 12 weeks. At the end of the 24-week study, serum and tissues from several organs, including the liver, adipose tissue, heart, kidneys, pancreas, and spleen, were collected. Liver tissue was used for histological, biochemical, and molecular analyses, as well as transcriptomic and metabolomic profiling (**Fig. 1A**). AATF expression was assessed across the collected tissues, confirming that AATF was silenced exclusively in the livers of AAV8-TBG-siAATF mice, compared to AAV8-TBG-siControl mice (**Fig. S1A** and **Fig. S1B**). Furthermore, AATF silencing was specific to hepatocytes, with no impact on non-parenchymal cells (**Fig. S1C**).

**Figure 1.**
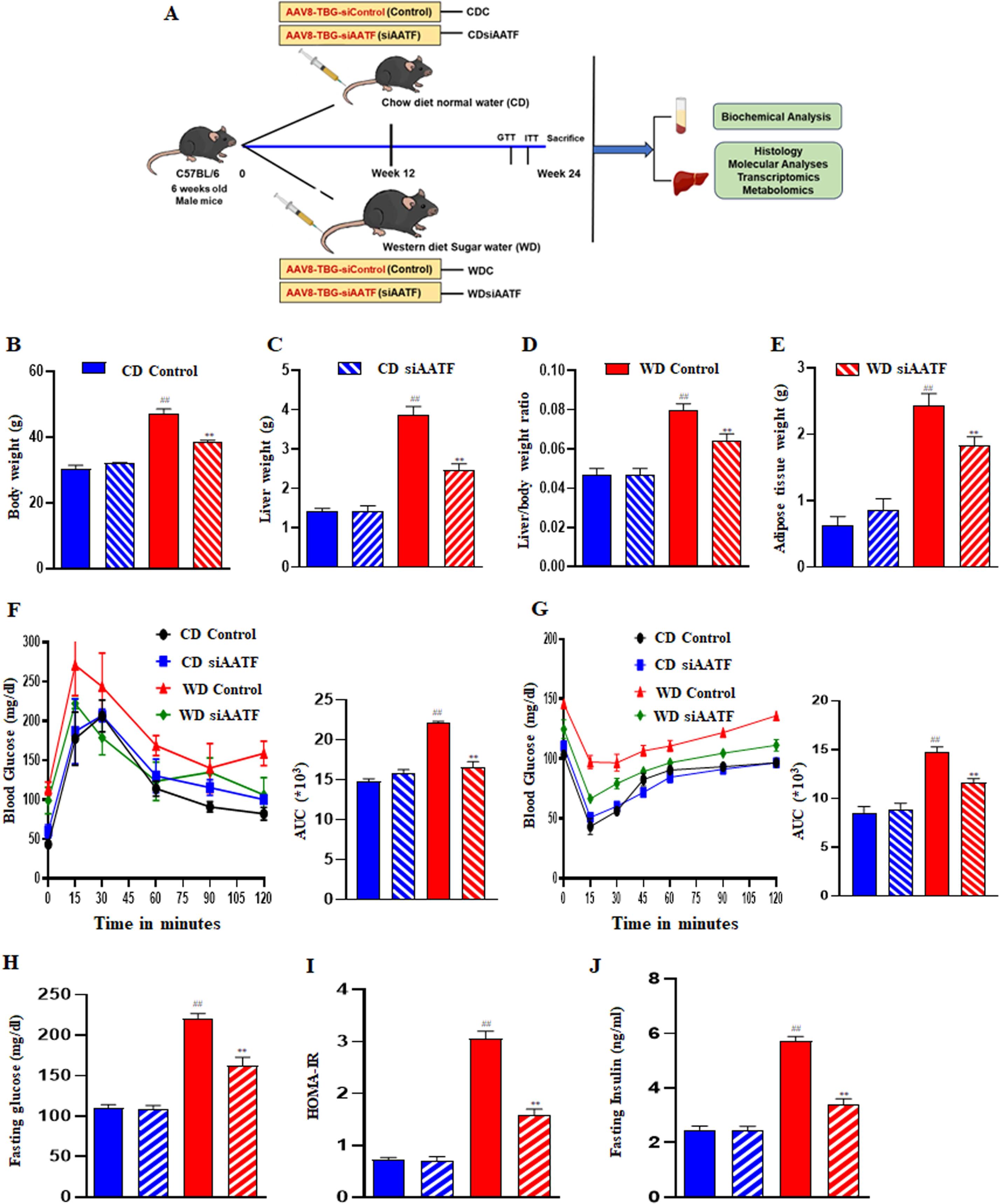
Hepatic silencing of AATF improves metabolic health in experimental MASH (A) Study design-C57Bl/6 mice were fed chow diet normal water (CD) or western diet sugar water (WD) for 12 weeks. Liver-specific AAV8-TBG-siControl (Control) or AAV8-TBG-siAATF (siAATF) were injected via the tail vein into mice and monitored for an additional 12 weeks. (B) Body weight, (C) liver weight, (D) liver-to-body weight ratio, and (E) adipose tissue weight were determined. Mean±SEM for 6-8 mice per group, ^##^p<0.001 compared to CD control; **p<0.001 compared to WD control, unpaired t-test. (F) Glucose tolerance test (G) Insulin tolerance test were performed. Mouse serum was collected after 24 weeks. (H) Fasting blood glucose, (I) fasting insulin, and (J) HOMA-IR were determined. Mean±SEM for 6-8 mice per group, ^##^p<0.001 compared to CD control; **p<0.001 compared to WD control, unpaired t-test. AAV; adeno-associated virus, TBG; tyrosine binding globulin, AATF; apoptosis antagonizing transcription factor, CD; chow diet, WD; western diet.

Silencing of hepatic AATF significantly reduced body weight, liver weight, liver-to-body ratio, and body fat mass in WD-fed mice, but did not affect CD mice (**Fig. 1B-1E, Fig. S2A**). Importantly, AAV8 administration had no adverse effects on the mice, as evidenced by stable bodyweight, normal behavior, and no mortality or abnormalities in major organs. Food intake was monitored throughout the study, and the cumulative calorie intake remained identical for all groups of mice **(Fig. S2B and S2C).** AATF silencing also improved glucose tolerance and insulin sensitivity in WD mice, as demonstrated by the glucose tolerance test (GTT) and the insulin tolerance test (ITT) (**Fig. 1F and 1G**). Notably, WD siAATF mice exhibited lower blood glucose and fasting insulin levels, along with a reduced homeostatic model assessment for insulin resistance (HOMA-IR) compared to WD control mice. These findings indicate AATF silencing ameliorates Western diet-induced impairments in glucose metabolism and insulin sensitivity (**Fig. 1H-1J**).

Furthermore, we also examined the effect of hepatic AATF silencing on liver injury by measuring serum levels of liver enzymes, including ALT, AST, total cholesterol, triglycerides, and LDL cholesterol. Consistently, WD siAATF mice showed reduced levels of liver enzymes compared to WD controls (**Fig. 2A-2E**). The macroscopic examination revealed that WD siAATF mice had a healthier liver morphology compared to WD controls (**Fig. 2F**). Histological analysis of liver sections showed decreased steatosis and substantial improvement in steatohepatitis upon hepatic AATF silencing compared to controls in WD mice (**Fig. 2G**). Similar results were obtained with Oil Red O staining of liver sections, where there was reduced intracellular lipid droplet accumulation in WD siAATF mice compared to WD controls (**Fig. 2H**). Consistent with these findings, histological scoring for steatosis, lobular inflammation, and hepatocyte ballooning was reduced in hepatic AATF-silenced WD mice, with an improvement in MASLD activity score compared to WD controls (**Fig. 2I**), indicating that hepatic AATF inhibition improves metabolic health by ameliorating liver injury and dysregulated lipid metabolism.

**Figure 2.**
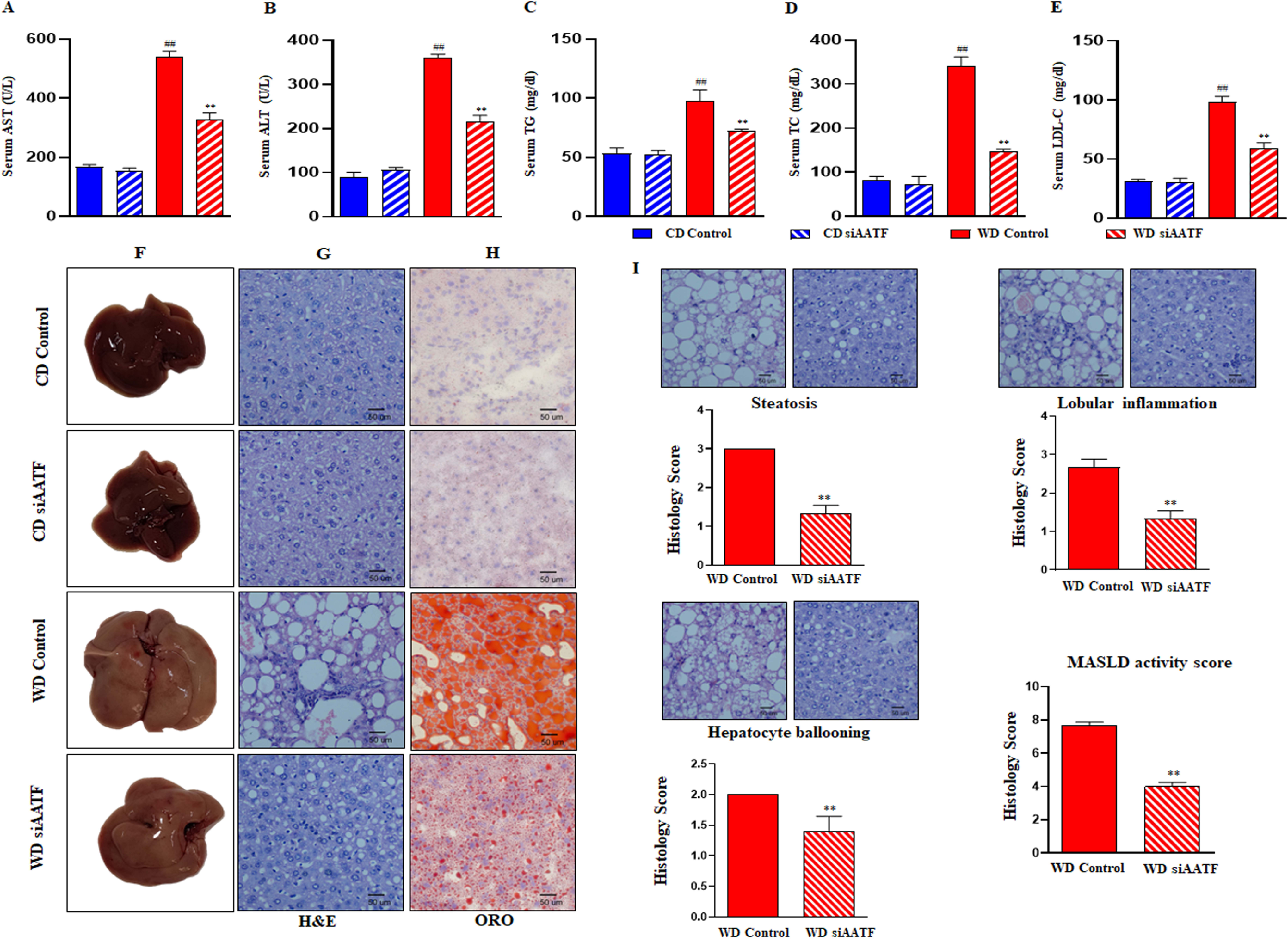
Hepatic loss of AATF improves liver injury and impaired lipid metabolism in MASH mice Mice were euthanized after 24 weeks, and blood was collected. Serum levels of (A) AST, (B) ALT, (C) total cholesterol, (D) triglycerides, and (E) LDL-cholesterol were measured. Mean±SEM for 6-8 mice per group, ^##^p<0.001 compared to CD control; **p<0.001 compared to WD control, unpaired t-test. Representative (F) gross liver images, (G) H&E staining, and (H) Oil Red O staining of the mouse liver tissues. n=6 mice per group. Scale bar 50 μm. (I) Histology scores were calculated for the liver sections of mice. Representative images and scoring of steatosis, lobular inflammation, hepatocyte ballooning, and MASLD activity score. Mean ±SEM for 6-8 mice per group. Scale bar 50 μm, unpaired t-test. **p<0.001 compared to WD control. HOMA-IR; homeostatic model assessment for insulin resistance, ALT, alanine transaminase; AST, aspartate transaminase; LDL; low-density lipoprotein, MASLD; metabolic dysfunction associated steatotic liver disease, CD; chow diet, WD; western diet.

### Effect of AATF silencing on hepatic transcriptomic and metabolomic profiles in MASH mice

To explore the liver-specific molecular pathways regulated by AATF in MASH, we performed RNA sequencing and untargeted metabolomic profiling of the liver tissues from all four groups. Comparisons were made between WDC (western diet-fed mice administered with AAV8-TBG-siControl) and CDC (chow diet-fed mice administered with AAV8-TBG-siControl), WDsiAATF (western diet-fed mice administered with AAV8-TBG-siAATF) and WDC (western diet-fed mice administered with AAV8-TBG-siControl), and CDsiAATF (chow diet-fed mice administered with AAV8-TBG-siAATF) and CDC (chow diet-fed mice administered with AAV8-TBG-siControl). The samples in transcriptomics analysis formed clear clusters between the WDsiAATF, WDC, and CD groups with good intra-group sample correlation (**Fig. 3A and 3B)**. Volcano plots showed a distribution of differentially altered genes in WDsiAATF vs. WDC (**Fig. 3C)**. Differential expression analysis showed 224 up and 144 downregulated genes in the WDsiAATF vs. WDC group (P value <0.05 and log 2-fold change≥2). In WDsiAATF mice, genes related to the anti-inflammatory response, such as neutrophilic granule protein (NGP), lactoferrin (LTF), and major urinary protein 1 (MUP1), which regulates gluconeogenesis and lipogenesis in the liver, were found to be upregulated. In contrast, genes like EPH receptor B2 (EPHB2) and suppression of tumorigenicity 18 (ST18), which play roles in liver fibrosis and inflammation, were downregulated. Additionally, the early growth response gene-2 (EGR2), which is crucial for the differentiation of monocytes into hepatic lipid-associated macrophages, and tubulin beta 2B class IIb (TUBB2B), involved in cholesterol biosynthesis, were also downregulated. GSEA analysis of the KEGG and GO databases showed suppressed extracellular matrix and collagen binding, immune response, activation of primary bile acid synthesis, and nitrogen metabolism (**Fig. 3D)**. The PLS-DA plot for hepatic metabolite profiling showed the differences between experimental groups and identified a significant separation of liver metabolites between the WDsiAATF and WDC groups, as well as the WDC and CDC groups (**Fig. 3E).** Univariate analysis identified 487 metabolites altered in the WD siAATF, of which 412 were upregulated, in contrast to 197 upregulated metabolites in WDC. Similarly, 867 metabolites were downregulated in WDC mice. In contrast, only 75 metabolites were downregulated in WDsiAATF mice, clearly indicating marked differences in metabolome profiling upon hepatic AATF silencing (Fig. 3F). Consistently, the volcano plots also showed a similar distribution of differentially altered metabolites in the WDsiAATF vs. WDC groups **(Fig. 3G).** No significant changes were observed in the hepatic transcriptomic and metabolomic profiles between CDsiAATF and CDC (**Fig. S3)**.

**Figure 3:**
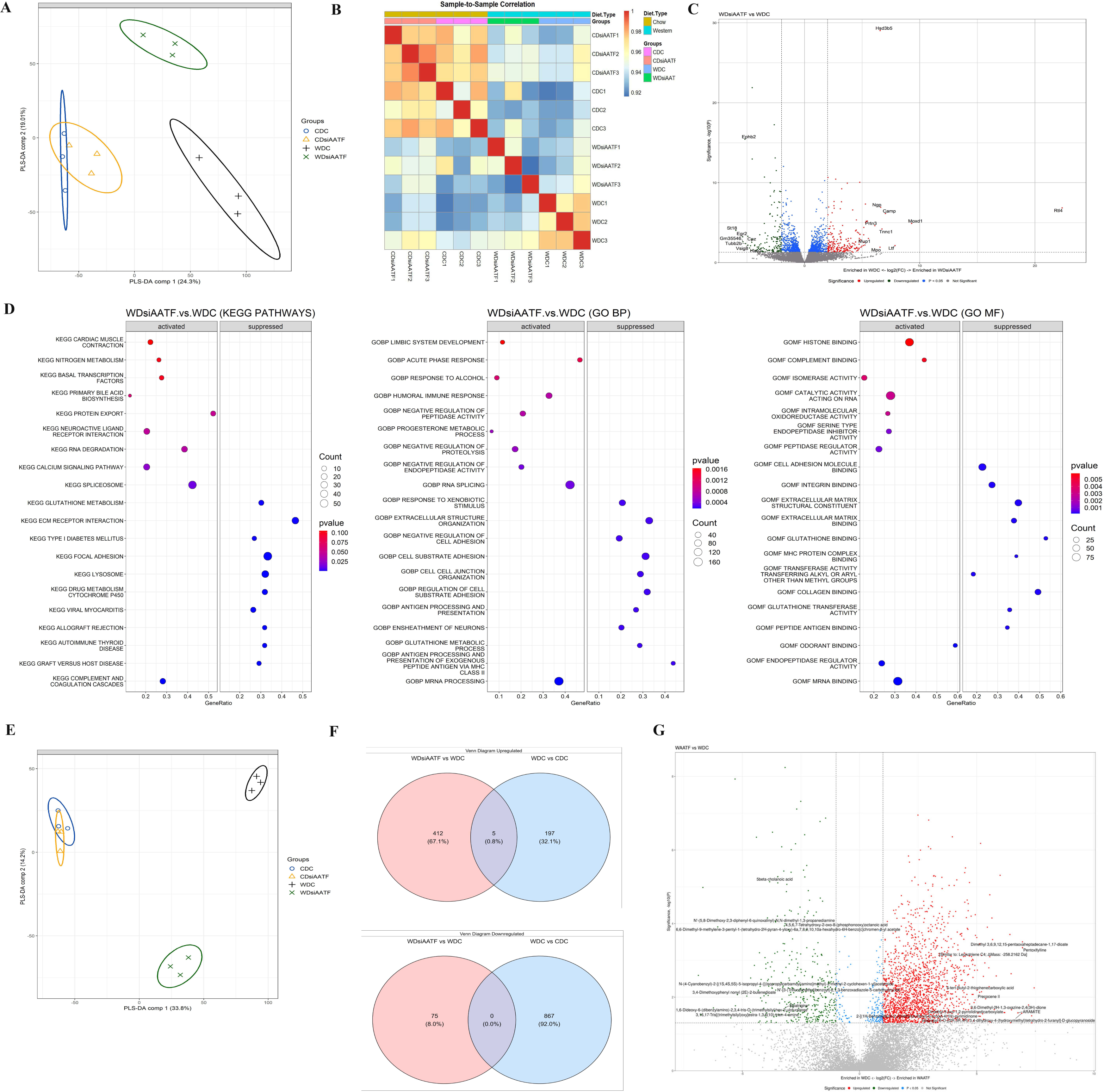
Effect of AATF silencing on hepatic transcriptomic and metabolic profiles in experimental MASH C57Bl/6 mice were fed chow diet normal water (CD) or western diet sugar water (WD) for 12 weeks. Liver-specific AAV8-TBG-siControl (Control) or AAV8-TBG-siAATF (siAATF) were injected via the tail vein into mice and monitored for an additional 12 weeks. After mice were euthanized, liver tissues were subjected to transcriptomic and untargeted metabolomic analyses. In transcriptomics, the PLS-DA plot (A) among the four groups shows a clear separation between the groups. (B) sample-sample correlation within the four groups. (C) Volcano plot for differentially expressed genes in WDsiAATF vs. WDC groups. The x-axis denotes fold change in expression (log2 scale), and the y-axis denotes adjusted p-value (negative log10 scale) for the analyzed genes in the dataset (each dot represents a single gene). Red and green dots represent genes significantly upregulated and downregulated, respectively. Blue dots (p<0.05) and grey dots represent nonsignificant genes. (DESeq2; cut-offs used: adjusted p-value < 0.05 and log fold change ≥ 2). (D) Top 10 up-and downregulated KEGG pathways, GO-BP, and GO-MF in WDsiAATF vs. WDC groups. In metabolomics, (E) PLS-DA plot among the four groups shows a clear group separation between the groups. (F) Venn diagram showing the up-and downregulated metabolites between the WDC vs. CDC and WDsiAATF vs. WDC. (G) Volcano plot for differential metabolites in WDsiAATF vs. WDC. Red and green dots represent metabolites significantly upregulated and downregulated, respectively. Blue dots (p<0.05) and grey dots represent nonsignificant metabolites. n=3. CD; chow diet, WD; western diet, AAV; adeno-associated virus, TBG; tyrosine binding globulin, AATF; apoptosis antagonizing transcription factor, PLS-DA; partial least squares discriminant analysis, WDC; western diet control, KEGG; Kyoto encyclopedia of genes and genomes, GO; gene ontology, BP; biological processes, MF; molecular function.

### Hepatic AATF loss alleviates cellular stress, inflammation, and fibrosis in experimental MASH

Cellular stress plays a prime role in the pathogenesis of MASH by promoting hepatic inflammation and fibrosis. Several factors contribute to cellular stress, stemming from lipid accumulation in hepatocytes, mitochondrial dysfunction, oxidative stress, and ER stress (30). In our investigation, we explored the impact of hepatic AATF silencing on cellular stress responses. Transcriptomics data showed downregulation of the ERK1/2 cascade, MAPK pathway, reactive oxygen species, PKA pathways, and apoptotic pathways in the WDsiAATF group (**Fig. 4A and 4B).** We subsequently validated the expression of key ER stress markers using qRT-PCR. Remarkably, in WDsiAATF mice, we found significant downregulation in the mRNA levels of CHOP, GRP78, PERK, and ATF4 compared to WDC (**Fig. 4C).**

**Figure 4.**
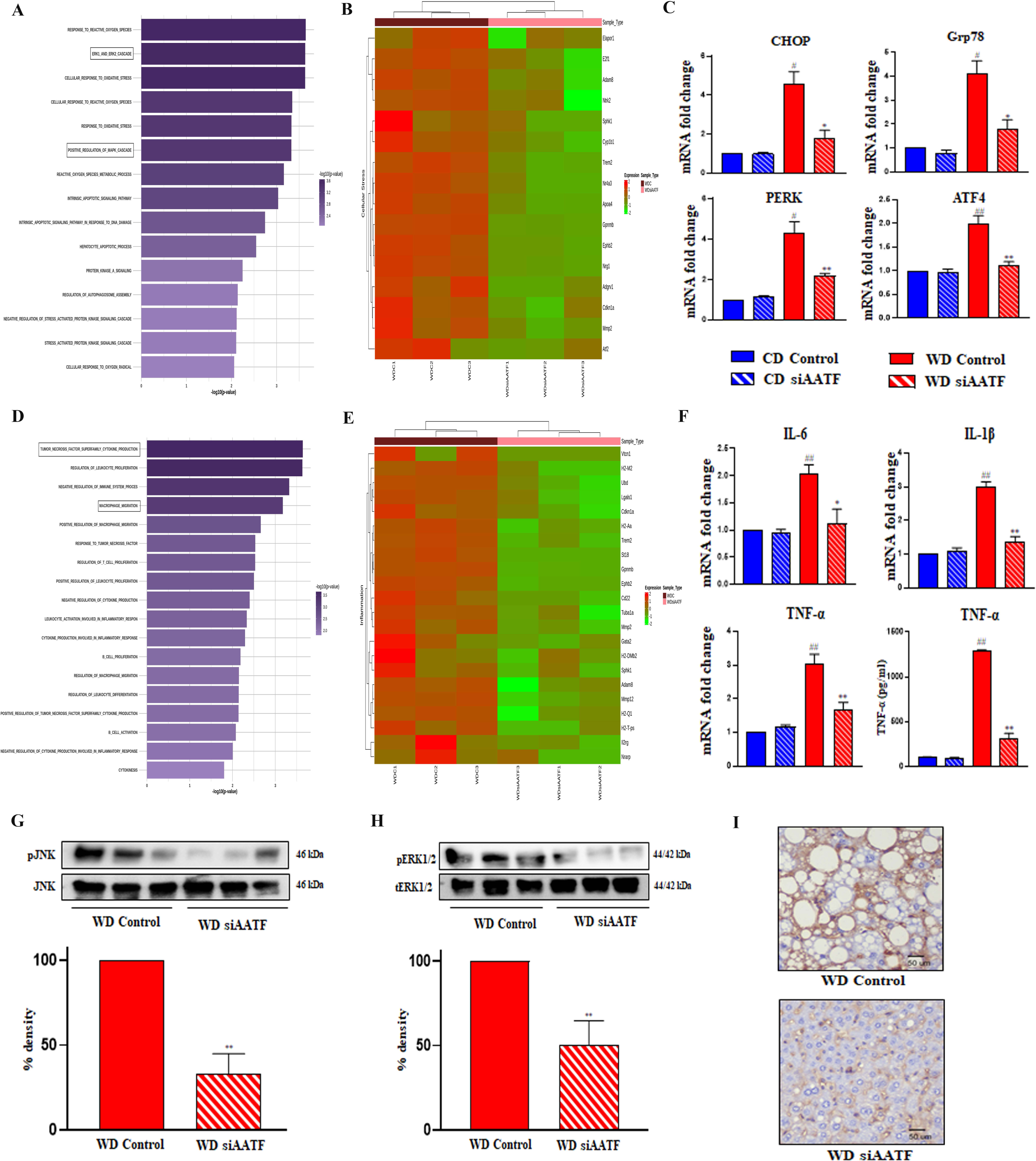
Effect of silencing hepatic AATF on cellular stress and inflammation in experimental MASH Transcriptomic analysis of liver tissues from mice fed with chow diet normal water (CD) or western diet sugar water (WD), administered with AAV8-TBG-siControl or AAV8-TBG-siAATF, n=3 (A) p-value plot of cellular stress-related pathways. (B) Heatmap of cellular stress-related genes. (C) hepatic mRNA expression of ER stress markers-CHOP, Grp78, PERK, and ATF4. Mean±SEM, n=4, unpaired t-test. (D) p-value plot of inflammation-related pathways. (E) Heatmap of inflammation-related genes. (F) hepatic mRNA expression of inflammatory markers-IL-1β, IL-6, and TNF-α, and serum TNF-α by ELISA. Liver tissue lysates were used for immunoblotting of inflammatory markers-(G) pJNK and (H) pERK1/2. (I) F4/80 staining, Scale bar 50 μm. Mean ±SEM, n=4, unpaired t-test. ^##^p<0.001 or ^#^p<0.05 compared to CD control; **p<0.001 or *p<0.05 compared to WD control. CD; chow diet, WD; western diet, AAV; adeno-associated virus, TBG; tyrosine binding globulin, AATF; apoptosis antagonizing transcription factor, CHOP; C/EBP homologous protein, GRP78; glucose-regulated protein 78, PERK; protein kinase R (PKR)-like endoplasmic reticulum kinase, ATF4; activating transcription factor 4, IL; interleukins, TNF-α; tumor necrosis factor-alpha, ELISA; enzyme-linked immunosorbent assay.

Furthermore, to elucidate the intricate crosstalk between cellular stress and inflammation, we hypothesized that hepatocellular stress triggers inflammatory signaling pathways, notably stress kinases, leading to the release of pro-inflammatory cytokines in WDC mice. Consequently, DEGs in WDsiAATF are attributed to the suppression of TNF-α superfamily cytokine production, macrophage migration, leukocyte proliferation, and differentiation (**Fig. 4D and 4E)**. Silencing of hepatic AATF in WD mice remarkably reduced inflammation, as evidenced by the reduced expression of pro-inflammatory cytokines IL-6, IL-1β, and TNF-α (**Fig. 4F**). Concurrently, phosphorylation of JNK and ERK1/2 was noticeably diminished in WDsiAATF mice compared to WDC (**Fig. 4G and 4H**). The immunostaining for the mouse macrophage marker F4/80 corroborated these findings, showing a marked reduction in response to AATF silencing in WD mice (**Fig. 4I**). These findings indicate that AATF is an important mediator of cellular stress-induced hepatic inflammation.

More interestingly, transcriptomic analysis of liver tissues from MASH mice revealed key genes associated with fibrosis, as evidenced by negative regulation of extracellular matrix organization, collagen metabolism, the TGF-β receptor signaling pathway, and fibroblast proliferation in WDsiAATF mice (**Fig. 5A and 5B)**. The further validation also confirmed reduced expression of α-SMA, Col1A1, Col3A1, and TGF-β, the key markers for liver fibrosis in the WDsiAATF group (**Fig. 5C**). Additionally, picrosirius red staining and hepatic stellate cell activation marked by desmin were attenuated in WDsiAATF mice, suggesting that AATF silencing reduces fibrogenic activation in WDC (**Fig. 5D and 5E).** Taken together, these results indicate the impact of hepatic silencing of AATF on key MASH signatures, including cellular stress, inflammation, and fibrosis.

**Figure 5.**
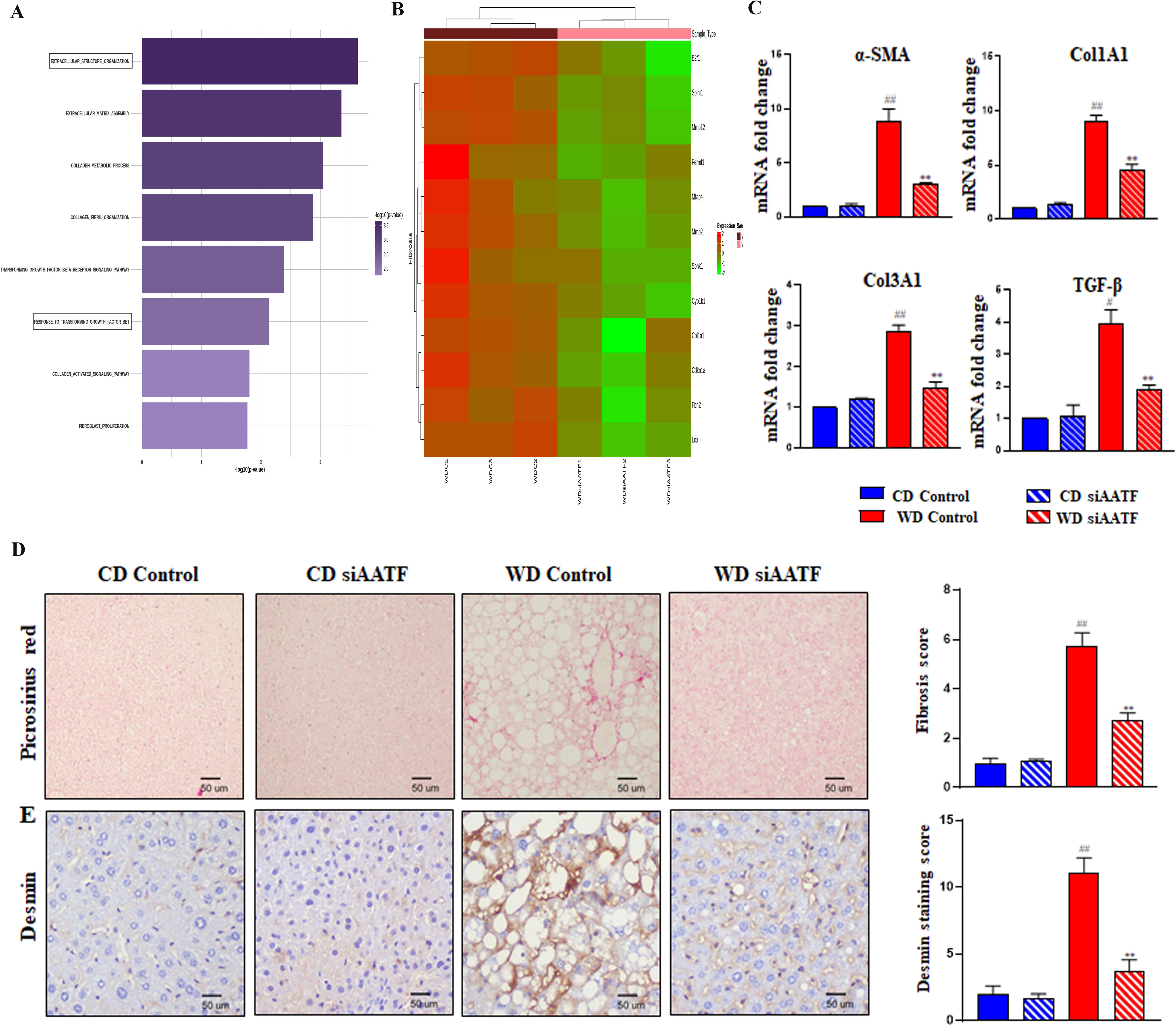
Silencing of hepatic AATF ameliorates liver fibrosis Transcriptomic analysis of liver tissues (n=3) showing (A) p-value plot of fibrosis-related pathways. (B) Heatmap of fibrosis-related genes. (C) Hepatic mRNA expression of Col1A1, Col3A1, α-SMA, and TGF-β quantified. Mean ±SEM, n=4, unpaired t-test. (D) Representative images of picrosirius red staining with fibrosis scoring. (E) Desmin immunostaining with scoring. Scale bar 50 μm. Mean ±SEM for 6-8 mice, unpaired t-test. ^##^p<0.001 or ^#^p<0.05 compared to CD control; **p<0.001 compared to WD control. Col; collagen, α-SMA; alpha-smooth muscle actin, TGF-β; transforming growth factor-beta. CD; chow diet, WD; western diet

### Hepatic loss of AATF regulates lipid metabolism and ameliorates metabolic dysfunction-associated steatohepatitis

Dysregulated lipid metabolism is the primary culprit in MASH pathogenesis, as it contributes to lipid accumulation, exacerbates insulin resistance, and triggers inflammation (31). This lipid accumulation results from excessive lipid influx, reduced fatty acid oxidation, and promotes de novo lipogenesis in the liver, ultimately leading to steatohepatitis (32). Differentially expressed genes (DEGs) from RNA sequencing in WDsiAATF were enriched in pathways such as the downregulation of ceramide and sphingolipids, the fatty acyl-CoA metabolic process, regulation of lipid localization, and transport (**Fig. 6A and 6B)**. Based on this observation, we further validated the genes involved in fatty acid synthesis, such as FASN, ACC1, ACC2, and SCD1. We found that lipogenic genes were significantly downregulated in WDsiAATF mice compared to WDC (**Fig. 6C)**.

**Figure 6.**
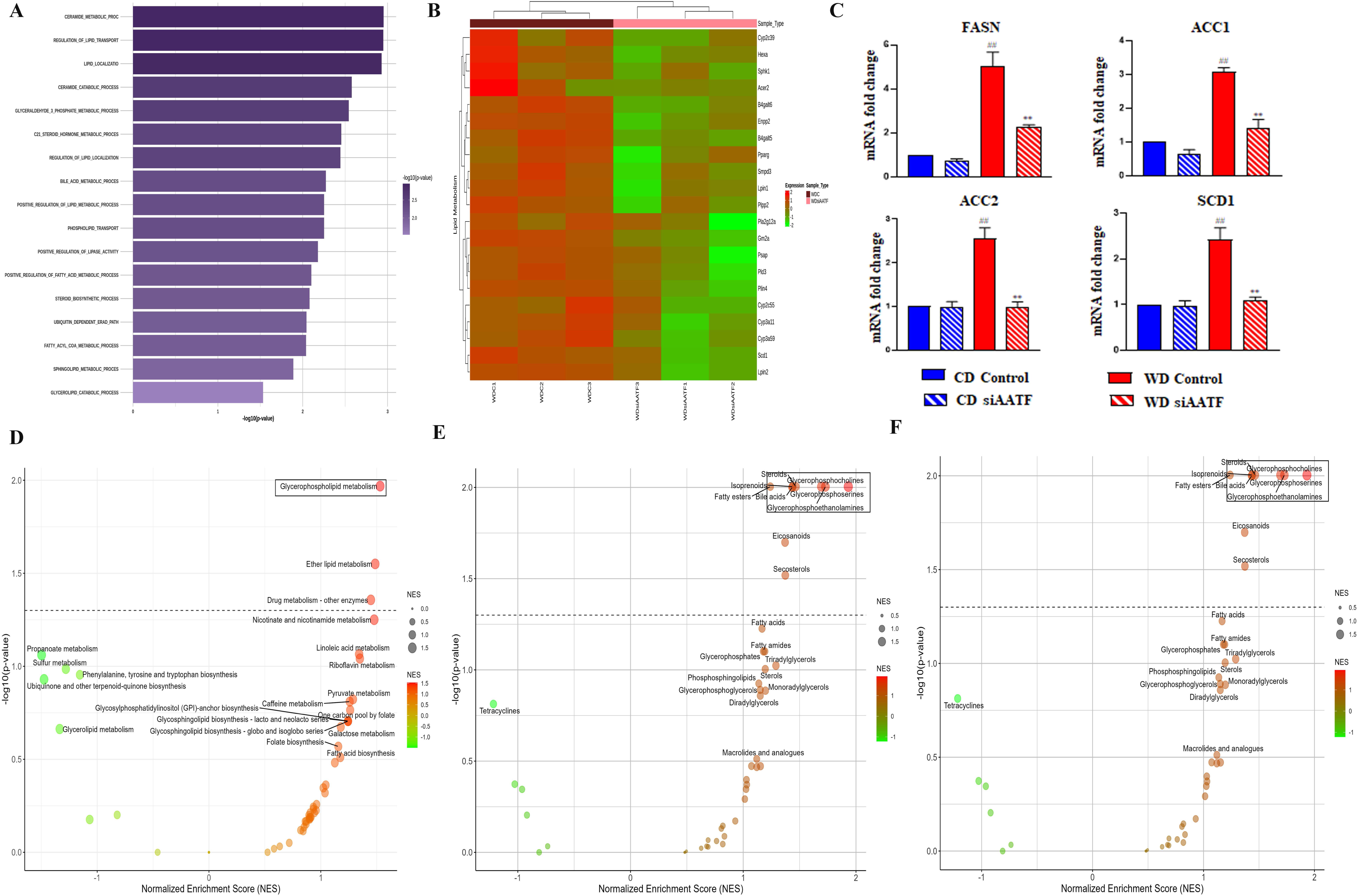
Effect of silencing hepatic AATF on lipid metabolism in MASH mice. Transcriptomic analysis of liver tissues (n=3) showing (A) p-value plot of lipid metabolism-related pathways. (B) Heatmap of lipid metabolism-related genes. (C) Hepatic mRNA expression of FASN, ACC1, ACC2, and SCD1 was evaluated. Mean ±SEM, n=4, unpaired t-test. Untargeted metabolomics was carried out in mouse liver tissues (n = 3). (D) Bubble plots showing the top 20 KEGG pathways enrichment, (E) lipid superclass, and (F) lipid subclass using the mummichog GSEA algorithm in WDC vs. WDsiAATF. P<0.05 was considered significant. CD; chow diet, WD; western diet, AAV; adeno-associated virus, TBG; tyrosine binding globulin, AATF; apoptosis antagonizing transcription factor, FASN; fatty acid synthase, ACC; acetyl-CoA carboxylase, SCD; stearoyl-coa 9-desaturase, KEGG; Kyoto encyclopedia of genes and genomes, GSEA; gene set enrichment analysis.

To comprehend the significance of the dysregulated metabolites for specific biological processes, a metabolic pathway analysis was conducted using MetaboAnalyst 6.0. The KEGG pathway enrichment analysis revealed that 74 pathways were enriched by the differentially expressed metabolites (DEMs) in the WD siAATF vs. WD control groups. Among these pathways, glycerophospholipid metabolism, ether lipid metabolism, nicotinamide metabolism, linoleic acid metabolism, riboflavin metabolism, and pyruvate metabolism were significantly upregulated upon hepatic silencing of AATF in MASH mice. In contrast, branched-chain amino acid biosynthesis, sulfur metabolism, and glycerolipid metabolism were downregulated **(Fig. 6D).** With the focus on only lipid-related pathways, DEMs were further enriched against the lipid superclass within the KEGG database, which unveiled the upregulation of glycerophospholipids (glycerophosphocholines, glycerophosphoserines, and glycerophosphoethanolamines), fatty esters, and eicosanoids in WDsiAATF mice **(Fig. 6E).** Delving deeper into lipid subclass analysis, PC, PS, PE, lysoPC, lysoPE, lysoPS, and vitamin D3 were elevated in WDsiAATF mice **(Fig. 6F).**

Glycerophospholipids are key structural elements of cell membranes and are involved in several cell signaling pathways (16). Notably, disruption in glycerophospholipid metabolism is known to compromise membrane stability and thereby contribute to the development of MASLD (17). Along the same lines, a network analysis of putative interacting proteins of the DEMs identified them as being enriched in the glycerophospholipid metabolism pathway in the KEGG database **(Fig. 7A)**. We further measured the relative abundance of different glycerophospholipids. We found that the levels of phosphocholine, PC, lysoPE, and lysoPC were significantly increased in WD siAATF mice compared to WD controls (**Fig. 7B**). This was further supported by the transcriptomic analysis, where the genes involved in glycerophospholipid metabolism were upregulated in WDsiAATF mice. (**Fig. 7C**). To gain deeper insights into the biological functions of the differentiated metabolites, a functional relationship network was constructed utilizing MetScape. The resulting network revealed that most of the metabolites were primarily involved in the metabolism of glycerophospholipids, linoleate, and phosphatidyl inositol (**Fig. S4**). Furthermore, we investigated the relationships between clinical variables related to MASH (anthropometry, liver enzymes, lipid profile, blood glucose, and insulin levels) and metabolite signatures using a debiases sparse partial correlation (DSPC) network based on MetaboAnalyst software (**Fig. 7D**). The correlation networks suggested that the majority of the metabolites belonging to glycerophospholipid metabolism showed strong positive interactions with each other and negative interactions with clinical variables, indicating a crucial role of glycerophospholipid metabolism in MASH pathogenesis. Collectively, these findings underscore the profound impact of AATF silencing on liver metabolites, particularly in the upregulation of glycerophospholipid biosynthesis.

**Figure 7.**
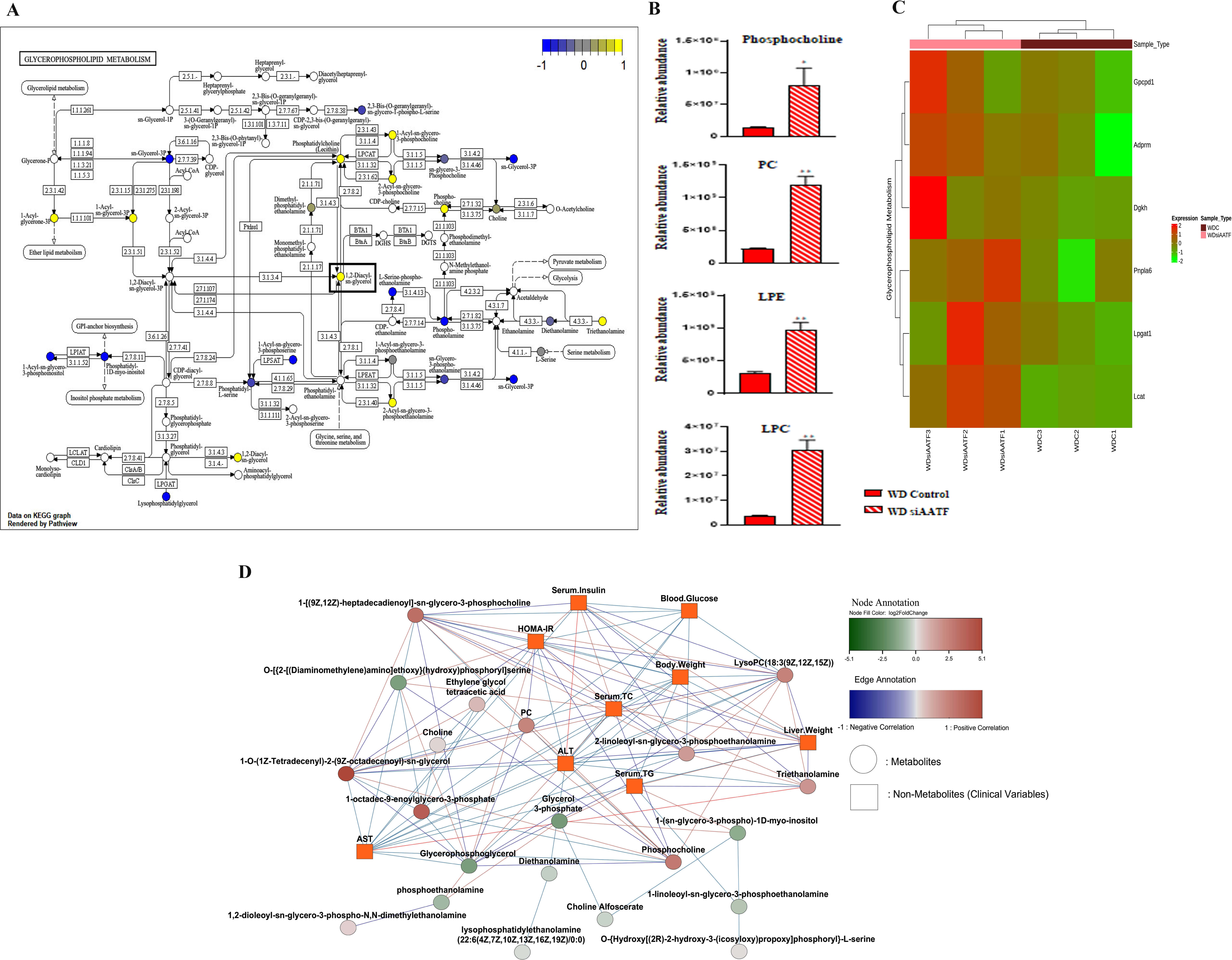
Metabolic networking of glycerophospholipid metabolism. Transcriptomics and untargeted metabolomics analyses were performed on mouse liver tissues (n = 3). Metabolites of glycerophospholipid metabolism were analyzed, and a functional correlation network analysis was performed. (A) The glycerophospholipid metabolism pathway, visualized in the Pathview network, highlights the dysregulated compounds (up-or downregulated) in WDsiAATF compared to WDC. (B) Relative abundance of phosphocholine, PC (phosphatidylcholine), LPE (lysophophatidylethanolamine), and LPC (lysophosphatidylcholine). **p<0.001 or *p<0.05 compared to WDC. (C) Heatmap of genes involved in glycerophospholipid metabolism. (D) Debiased sparse partial correlation network plot visualizing the major relationships of glycerophospholipid metabolic alterations with the clinical variables. Statistical significance was analyzed by a student’s t-test. P<0.05 was considered significant. CD; chow diet, WD; western diet, AATF; apoptosis antagonizing transcription factor.

The process of beta-oxidation of fatty acids plays a crucial role in the development of MASH. When beta-oxidation pathways are disrupted, it leads to the accumulation of lipids in the liver, increased oxidative stress, and inflammation, all of which contribute to disease progression (31). In contrast, alpha-linolenic acid (ALA), an essential omega-3 (n-3) polyunsaturated fatty acid (PUFA), provides substantial antioxidant benefits, as well as anti-inflammatory and lipid-lowering effects (33). Along similar lines, our studies revealed the upregulation of metabolites involved in the mitochondrial beta-oxidation pathway and alpha-linolenic acid metabolism alongside the biosynthesis of glycerophospholipids in WDsiAATF through pathway enrichment analysis using the SMPB database (**Fig. 8A**). Consistently, genes involved in fatty acid oxidation were upregulated in WDsiAATF mice (**Fig. 8B)**. Based on this, we hypothesized whether silencing AATF enhances beta-oxidation of fatty acids, thereby improving MASH. Supporting this, our results showed a significant upregulation of key genes involved in fatty acid oxidation, including carnitine palmitoyltransferase-1 alpha (CPT1α), acyl-CoA oxidase 1 (ACOX1), peroxisome proliferator-activated receptor gamma coactivator-1 alpha (PGC1α), and acyl-CoA synthetase long-chain family member 1 (ACSL1) in WD siAATF mice compared to the controls (**Fig. 8C**). Furthermore, to understand the possible interacting partners of AATF in regulating MASH progression, we performed an integrative analysis of AATF interacting proteins with the transcriptomic signature of the WDsiAATF group using the STRING and BioGRID databases (**Fig. S5)**. The protein-protein interaction network revealed that AATF is involved in the positive regulation of fatty acid metabolic processes through CD74 and MAPK9; however, further validation is necessary to elucidate the signaling pathway.

**Figure 8.**
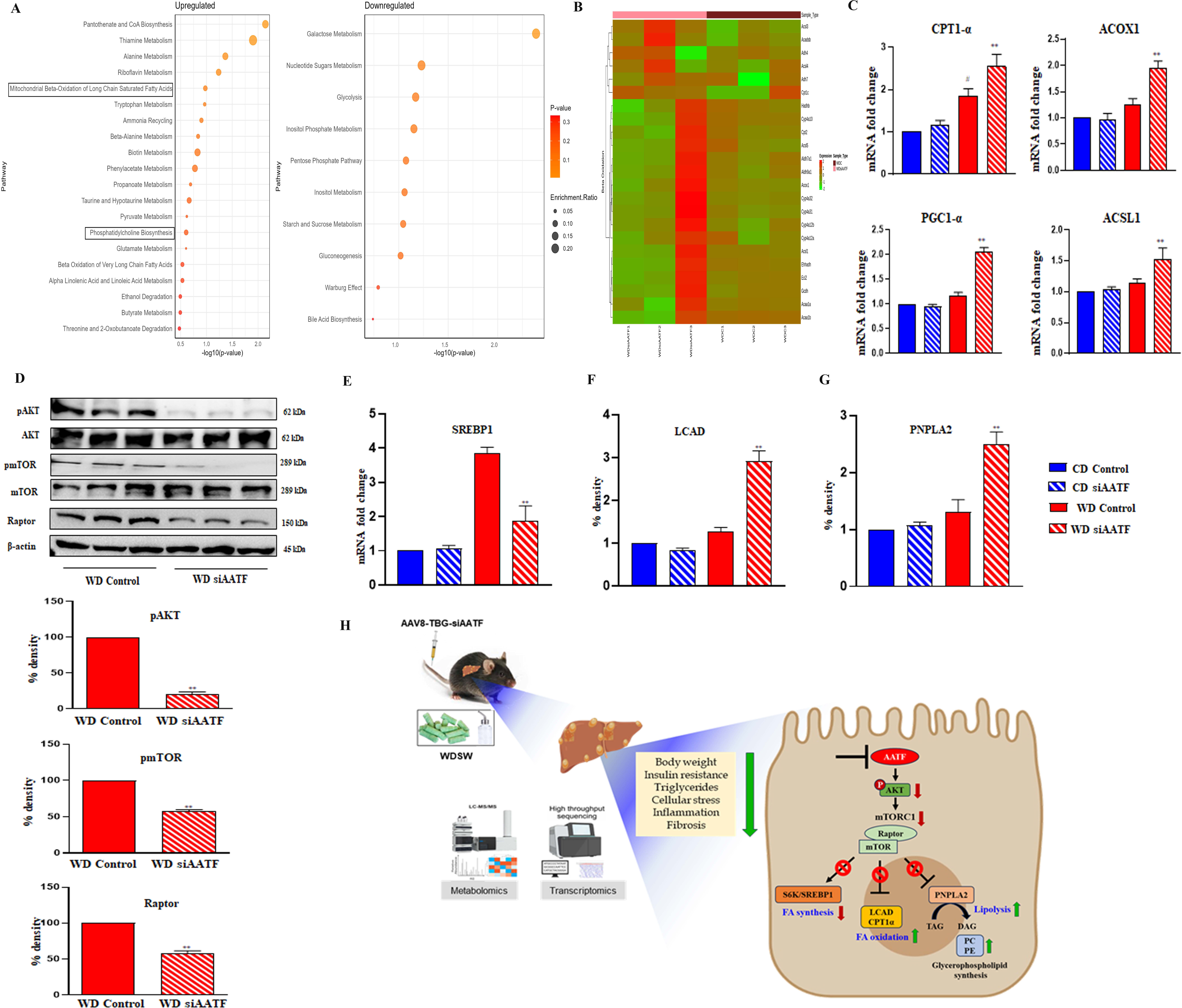
Hepatic loss of AATF alters lipid metabolism via AKT-mTORC1 signaling pathways in MASH mice Transcriptomics and untargeted metabolomics analyses were performed on mouse liver tissues (n = 3). Pathway analysis and enrichment analysis were performed on both datasets. An enrichment analysis based on the Small Molecule Pathway Database (SMPDB) was performed on metabolites. (A) upregulated and downregulated pathways. (B) Heatmap of genes involved in fatty acid oxidation. (C) hepatic mRNA expression of CPT-1α, ACOX1, PGC-1α, and ACSL1. (D) Liver tissue lysates were used for immunoblotting of pAKT (Thr308), pmTOR, and Raptor. Hepatic mRNA expression of (E) SREBP1, (F) LCAD, and (G) PNPLA2. Mean±SEM, n=4, unpaired t-test, **P < 0.001 or *P < 0.05 compared to WDC. (H) Schematic representation of the effect of hepatic AATF silencing in experimental MASH. ACOX1; Acyl-CoA Oxidase 1, PGC1; Peroxisome proliferator-activated receptor gamma coactivator-1alpha, ACSL1; acyl-CoA synthetase long-chain family member 1, AKT; protein kinase B, mTOR; mammalian target of rapamycin, SREBP1; sterol regulatory element-binding protein 1, LCAD; long-chain acyl-CoA dehydrogenase, PNPLA2; patatin-like phospholipase domain-containing protein 2.

### Molecular mechanisms underlying hepatic AATF-driven MASH progression

mTORC1 (mammalian target of rapamycin complex 1) is a central regulator of cell growth and metabolism that contributes to the pathogenesis of MASH through its effects on lipid metabolism (34–36). In MASH, the activation of the AKT (protein kinase B) and mTORC1 signaling pathways is often dysregulated, contributing to the disease progression (37). We hypothesized that AATF silencing inactivates the AKT-mTORC1 signaling pathway, thereby mobilizing hepatic lipid accumulation. Accordingly, we found that phosphorylation of AKT at Thr308 was inhibited in WDsiAATF mice compared to the WDC mice. Following this, activation of mTORC1, represented by the phosphorylation of mTOR and Raptor, was downregulated in WDsiAATF mice (**Fig. 8D)**. Further, SREBP1 levels were downregulated in WDsiAATF mice, supporting the mTORC1-mediated lipogenesis (**Fig. 8E**). Previous studies have shown that mTORC1 inhibits lipolysis and β-oxidation of fatty acids (38,39). In our study, we found that inactivation of mTORC1 via AATF activates long-chain acyl-CoA dehydrogenase (LCAD), an enzyme crucial for the initiation of β-oxidation (**Fig. 8F**). Interestingly, PNPLA2 (Patatin-Like Phospholipase Domain-Containing Protein 2), an enzyme involved in lipolysis, particularly breakdown of triglycerides, was upregulated in WDsiAATF mice compared to WDC (**Fig. 8G**). PNPLA2 is responsible for the conversion of triglycerides to diacylglycerol and free fatty acids, which further undergo glycerophospholipid synthesis and β-oxidation, respectively (40). Collectively, hepatic silencing of AATF ameliorates MASH by enhancing glycerophospholipid biosynthesis and β-oxidation of fatty acids via inhibition of AKT-mTORC1 signaling pathways (**Fig. 8H**).

## Discussion

MASLD is a multifactorial condition influenced by various molecular drivers (5). Dysregulation of lipid metabolism is a hallmark of MASLD. The increased influx of free fatty acids (FFAs) from adipose tissue and dietary sources, combined with impaired fatty acid oxidation and the export of triglycerides from the liver, contributes to hepatic steatosis (11). Insulin resistance in hepatocytes leads to impaired suppression of gluconeogenesis and increased lipogenesis, promoting hepatic lipid accumulation (31). Several factors associated with metabolic dysfunction, including lipid accumulation, oxidative stress, and insulin resistance, can trigger ER stress in hepatocytes (30). Inflammatory pathways play a crucial role in the progression of MASLD from simple steatosis to MASH. Activation of inflammatory signaling pathways leads to the production of pro-inflammatory cytokines and chemokines, contributing to hepatocyte injury and fibrosis (41). Thus, identifying and understanding the molecular drivers involved in MASH pathogenesis is essential for developing targeted therapeutic strategies aimed at preventing disease progression and improving clinical outcomes for patients with MASLD. The evolving understanding of the disease underscores the urgent need for new therapeutic targets. The complex pathophysiology of MASH involves multiple metabolic, inflammatory, and fibrotic pathways, making it essential to identify novel targets that can address these different aspects. Advanced molecular profiling techniques, including transcriptomics, proteomics, metabolomics, lipidomics, and epigenomics, can provide valuable insights into the underlying mechanisms driving MASH and guide the development of personalized treatment approaches.

AATF has been implicated in the pathogenesis of several cancers due to its role in modulating apoptotic pathways (20). We, for the first time, elucidated how TNF-α upregulates the expression of AATF via SREBP-1, promoting hepatocarcinogenesis (42). Further studies revealed the role of AATF in impeding HCC angiogenesis by downregulating PEDF (26). Recently, we investigated the role of AATF in MASH-HCC and showed that TACE inhibition by Marimastat prevents AATF-mediated inflammation, fibrosis, and oncogenesis (28). Building on these experimental findings, we sought to understand the role of AATF in MASH. In the current study, we explored for the first time how AATF regulates lipid metabolism and its implications in the development and progression of MASH. We employed a preclinical murine model of MASH comprising C57BL/6 mice that were fed a standard chow diet, normal water, or western diet sugar water. This model effectively mimics the features of metabolic syndrome and MASH, including the progression of fibrosis. The mice develop fatty liver within 12 weeks of being fed with a western diet, sugar water, progressing to MASH and fibrosis by 24 weeks. This spectrum mirrors the dietary pattern, systemic milieu, and metabolic and histological characteristics of MASH, demonstrating the activation of key cellular pathways associated with human disease. This approach overcomes the constraint of the previously employed DIAMOND mouse model, which was an isogenic cross between C57BL/6J and 129S1/SvImJ mouse strains (29). Although the DIAMOND model closely mimics the human disease characteristics, the genetic instability and associated inconsistency in phenotypic outcomes render it less accepted. In the current study, we assessed the impact of liver-specific silencing of AATF on hepatic lipid accumulation in experimental MASH. Of note, while siRNA was chosen for gene silencing, the approach employed in this study closely resembles the characteristics of shRNA, particularly in terms of its stability. The technology developed by Applied Biological Materials Inc. (ABM) utilizes a convergent promoter vector system, which allows siRNA to be expressed in both directions— a feature typical of shRNA. Additionally, the AAV8 vector used for gene delivery offers tissue-specific targeting, providing stability and sustained gene expression. We examined how this silencing affected the genes, proteins, and metabolites associated with the biological processes involved in the development and progression of MASH. The silencing of hepatic AATF showed a significant reduction in body weight and liver weight in WDSW mice and did not affect CDNW mice. Insulin resistance was notably improved, and we also observed substantial improvement in both histological and biochemical parameters in WDsiAATF mice compared to WDC. In line with these findings, liver transcriptomics analysis and on-bench validations demonstrated that the loss of AATF reduced cellular stress responses, leading to decreased production of inflammatory cytokines and fibrogenesis in WDSW mice. These results underscore the potential role of AATF in the progression of MASLD and set the groundwork for more detailed molecular investigations.

This is the first study to conduct both hepatic transcriptomic and metabolomic profiling in mice with AATF silenced. Since the precise mechanism(s) by which the loss of AATF alleviates MASH are unclear, we concentrated on detecting changes in liver metabolites in the experimental MASH group with AATF silencing compared to the control group using untargeted metabolomics, which offers a comprehensive overview of the entire metabolome. Based on transcriptomic analysis and our key interest in elucidating the role of AATF in lipid metabolism, we found it intriguing that the leading unregulated pathways among the 74 pathways enriched by DEMs in the WD siAATF compared to the WD control group included glycerophospholipid metabolism, ether lipid metabolism, nicotinamide metabolism, linoleic acid metabolism, riboflavin metabolism, and pyruvate metabolism. Notably, there was a marked increase in glycerophospholipids, particularly PC, PS, and PE, in the WD siAATF mice. These subclasses of lipids are implicated in MASLD progression, where dysregulation of these glycerophospholipids is linked with insulin resistance, inflammation, and fibrosis, which are the hallmarks of MASLD (43). Additionally, their breakdown products, like LysoPC, LysoPS, and LysoPE, were also increased. The metabolite-metabolite interaction network highlighted the potential functional relationships of glycerophospholipid with linoleate metabolism. It is noteworthy that ALA, an essential omega-3 fatty acid, is primarily attributed to its anti-inflammatory, antioxidant, and lipid-lowering properties in MASLD (44). ALA can influence lipid metabolism by modulating the expression of genes involved in fatty acid oxidation and lipid synthesis. It also promotes the activation of peroxisome proliferator-activated receptors (PPARs), which play a crucial role in lipid metabolism (45).

Previous studies have also reported alterations in phospholipid content in the liver biopsies of MASH patients, suggesting that impaired phospholipid synthesis is associated with disease progression (14,15). The low hepatic PC levels and the altered hepatic PC to PE ratio appear to play significant roles in the development of MASLD (46). However, the pathophysiology of these lipid-induced processes remains not fully understood. Moreover, low PC levels have previously been shown to activate SREBP1 (47). Essential phospholipids (EPLs) rich in PC are a widely used treatment option for fatty liver disease. There is substantial and consistent clinical evidence indicating that treatment with EPLs leads to the regression of steatosis (48). Studies show that phosphatidylcholine metabolism intersects with various metabolic pathways, and optimizing PC levels could potentially help regulate lipid metabolism and insulin sensitivity, thereby mitigating the progression of MASLD (49). Additionally, PC may possess antioxidant properties, which could counteract oxidative damage in the liver and protect against disease progression (50). In addition to various lipid classes, the liver metabolite profile in WDsiAATF mice showed decreased branched-chain amino acid metabolism and sulfur metabolism involved in the MASH pathogenesis, in contrast to increased pyruvate metabolism, NAD, and FAD metabolism, as well as one-carbon metabolic pathways. The effect of AATF silencing on non-lipid metabolic pathways reveals new dimensions of AATF’s functional significance in MASLD (data not shown; manuscript under preparation).

Another intriguing observation in the current study using pathway enrichment analysis is the upregulation of mitochondrial beta-oxidation of the fatty acid pathway in WDsiAATF mice. Fatty acid oxidation (FAO) helps to efficiently metabolize fatty acids, and thereby enhanced beta-oxidation may alleviate inflammation and oxidative stress in the liver, which are key drivers of MASLD progression (51). Several genes, such as CPT1α, PPARs, and ACOX1, encode enzymes involved in FAO, and henceforth, upregulated expression of these genes would enhance fatty acid oxidation capacity, potentially improving metabolic health (52,53). Consistent with this, our study also showed the upregulation of FAO genes in the liver tissue of WDsiAATF compared to WDC. The development of hepatic steatosis is primarily associated with the excessive accumulation of triglycerides in hepatocytes. Our research suggests that loss of hepatic AATF may redirect the liver’s metabolism from triglyceride synthesis to the production of diacylglycerol (DAG) and, subsequently, glycerophospholipids, helping to manage or decrease fat accumulation in the liver.

Furthermore, increased fatty acid oxidation could mobilize the stored triglycerides, improving metabolic balance, reducing oxidative stress, and potentially alleviating MASH. An in-depth, targeted lipidomics approach coupled with triglyceride profiling will yield definitive insights, which is the central focus of our forthcoming research. Interestingly, the protein-protein interaction network of AATF-interacting proteins, in conjunction with the transcriptomic profile of WDsiAATF, revealed CD74 as a possible link between AATF and lipid metabolism. Supporting this, studies by Li et al. have shown that CD74 levels were positively correlated with triglyceride levels in human atherosclerotic lesions (54).

Studies by Ishigaki et al. have demonstrated that AATF activates AKT in response to cellular stress, thereby promoting cell survival (55). AKT phosphorylated at Thr308 activates mTORC1, which particularly plays a dominant role in promoting lipogenesis and inhibiting lipid breakdown (56, 57). Thus, we evaluated the effect of the AATF-AKT-mTOR signaling pathway on hepatic lipid metabolism. AATF silencing effectively inactivated mTORC1 activation by AKT, further affecting the lipogenic genes such as SREBP1, FASN, and ACC. The inactivated mTORC1 complex led to the upregulation of LCAD, a key enzyme required for initiating the β-oxidation of fatty acids. Moreover, mTORC1 suppresses PNPLA2, a key lipase involved in the lipolysis of triglyceride to diacylglycerol and free fatty acids (58). The stored fat is mobilized in the form of diacylglycerol, a precursor for glycerophospholipids and free fatty acids, which undergo β-oxidation for energy production. In our study, AATF silencing increased PNPLA2 levels, as reflected by decreased triglyceride levels and upregulated glycerophospholipid synthesis and β-oxidation in WDsiAATF mice.

Despite extensive studies on MASLD, understanding the molecular drivers of lipid metabolism, insulin resistance, oxidative stress, and inflammation is essential for addressing its pathogenesis. Although knockout models remain the gold standard for functional validation, we utilized AAV8-mediated, hepatocyte-specific siRNA delivery to achieve temporal and liver-restricted silencing of AATF. This approach avoids developmental compensations inherent to germline knockouts and offers translational relevance, mimicking therapeutic gene silencing strategies. The use of the TBG promoter ensured hepatocyte specificity, minimizing off-target effects in non-parenchymal cells. Given AATF’s pleiotropic functions, a complete knockout may result in systemic effects not confined to liver pathology. Importantly, AAV8 vectors are widely used in translational gene therapy platforms, providing a clinically relevant proof of concept. While we acknowledge that future studies using hepatocyte-specific AATF knockout mice will be essential to validate the long-term impact and specificity of our findings fully, our results provide compelling evidence for the functional role of AATF in MASH and establish a foundation for therapeutic targeting. Another promising alternative is the development of small-molecule inhibitors targeting AATF, offering a more practical and sustained therapeutic approach. Identifying and optimizing these inhibitors will be critical for advancing AATF-targeted therapies for MASH and other metabolic diseases.

In conclusion, this study identifies AATF as a novel transcriptional regulator of lipid metabolism in the context of MASH. It elucidates a previously unknown AKT– mTORC1 axis, which is modulated by AATF, in hepatocytes. Our use of liver-specific AAV8-mediated siRNA delivery enabled sustained and targeted gene silencing, representing a translationally relevant model for future therapeutic development. Furthermore, the application of integrated transcriptomics and metabolomics provides a systems-level understanding of how AATF silencing reprograms lipid metabolism, particularly through the upregulation of glycerophospholipid biosynthesis and fatty acid oxidation. These findings open new avenues for precision-targeted treatments and highlight AATF as a promising candidate for pharmacological inhibition in MASH.

## Declarations

## Ethics approval and consent to participate

Not applicable

## Consent for publication

Not applicable

## Availability of data and materials

Data presented herein are available from the corresponding author upon request. RNA sequencing data have been uploaded to the GEO database (Accession number: GSE280320)

## Competing interests

The authors declare that there are no conflicts of interest.

## Funding

This study was supported in whole or in part by the SERB-POWER grant (SPG/2021/002524) and Ramalingaswami Re-entry Fellowship from the Department of Biotechnology (BT/RLF/Reentry/58/2017) to DPK, and Senior Research Fellowship (SRF) from ICMR to ANS.

### Authors’ contributions

Conceptualization: ANS, DPK; Methodology: ANS, DS, MM, PA, DPK; Investigation: ANS, PKS, DS, SPS, SMN, DPK; Formal Analysis: ANS, DS, MM, GR, PKS, DPK; Visualization: ANS, DPK; Resources: ANS, MM, GR, SPS, SMN, PKS, DPK; Supervision: ANS, DPK; Funding Acquisition: DPK; Writing-original draft: ANS, DPK; Writing-Review and Editing: All authors.

## Acknowledgements

RNA sequencing was carried out at Molsys Pvt. Ltd. (Molsys Scientific). We thank the DBT-SAHAJ National Facility at Rajiv Gandhi Centre for Biotechnology (RGCB) for their work on metabolomics. We also thank TheraCUES Innovations Pvt Ltd. for assisting with bioinformatic analysis of the transcriptomics and untargeted metabolomics assay data.

The authors also acknowledge funding support from the Department of Biotechnology-Boost to University Interdisciplinary Life Science Departments for Education and Research program [DBT-BUILDER: BT/INF/22/SP43045/2021] and the Department of Science and Technology–Promotion of University Research and Scientific Excellence [DST-PURSE: SR/PURSE/2021/81(c)].

---

**Supplemental Figure 1:** (A) Tissue distribution of Control and siAATF in the experimental mice. (B) liver-specific, and (C) hepatocyte-specific AATF silencing as determined by qRT-PCR and western blot, respectively. Mean±SEM for n=4 mice per group, **p<0.001 compared to AAV8 control, unpaired t-test.

**Supplemental Figure 2:** (A) Weekly measurement of body weight from week 12 to week 24, followed by the administration of AAV8-TBG-siControl (Control) and AAV8-TBG-siAATF (siAATF) at week 12 to the chow diet normal water (CD) and western diet sugar water (WD) fed mice. (B) Food intake and (C) Calorie intake in mice. Mean±SEM for 6-8 mice per group, one-way ANOVA.

**Supplemental Figure 3:** (A) Volcano plot presentation of differentially expressed genes between liver samples of mice from WDC and CDC. (B) Volcano plot presentation of differentially expressed genes between liver samples of mice from CDsiAATF and CDC. (C) Volcano plot presentation of differential metabolites from liver samples of mice from WDC and CDC. (D) Volcano plot presentation of differential metabolites from liver samples of mice from CDsiAATF and CDC. X-axis denotes fold change in expression (log2 scale), y-axis denotes adjusted p-value (negative log10 scale) for the analyzed genes or metabolites in the data set (each dot represents a single gene or metabolite). Red and green dots represent genes or metabolites significantly upregulated and downregulated, respectively. Blue dots (p<0.05) and grey dots represent nonsignificant genes or metabolites. (DESeq2; cut-offs used: adjusted p-value < 0.05 and log fold change ≥ 2).

**Supplemental Figure 4:** Metabolite functional interaction network of glycerophospholipid metabolism pathway in WDsiAATF and WDC.

**Supplemental Figure 5:** Protein-protein interaction of AATF interacting proteins with the lipid metabolism-related proteins was determined using BioGRID and STRING databases and visualized in Cytoscape.

**Supplemental Table 1:** List of primer sequences used in qRT-PCR in the study.

## Notes

### Competing Interest Statement

The authors have declared no competing interest.

### Summary of Updates

In this version, the title of the manuscript has been modified and study novelty has been highlighted.

