## Supplementary data for "Hepatic loss of AATF attenuates MASH by suppressing AKT–mTORC1 signaling and reprogramming lipid metabolism"

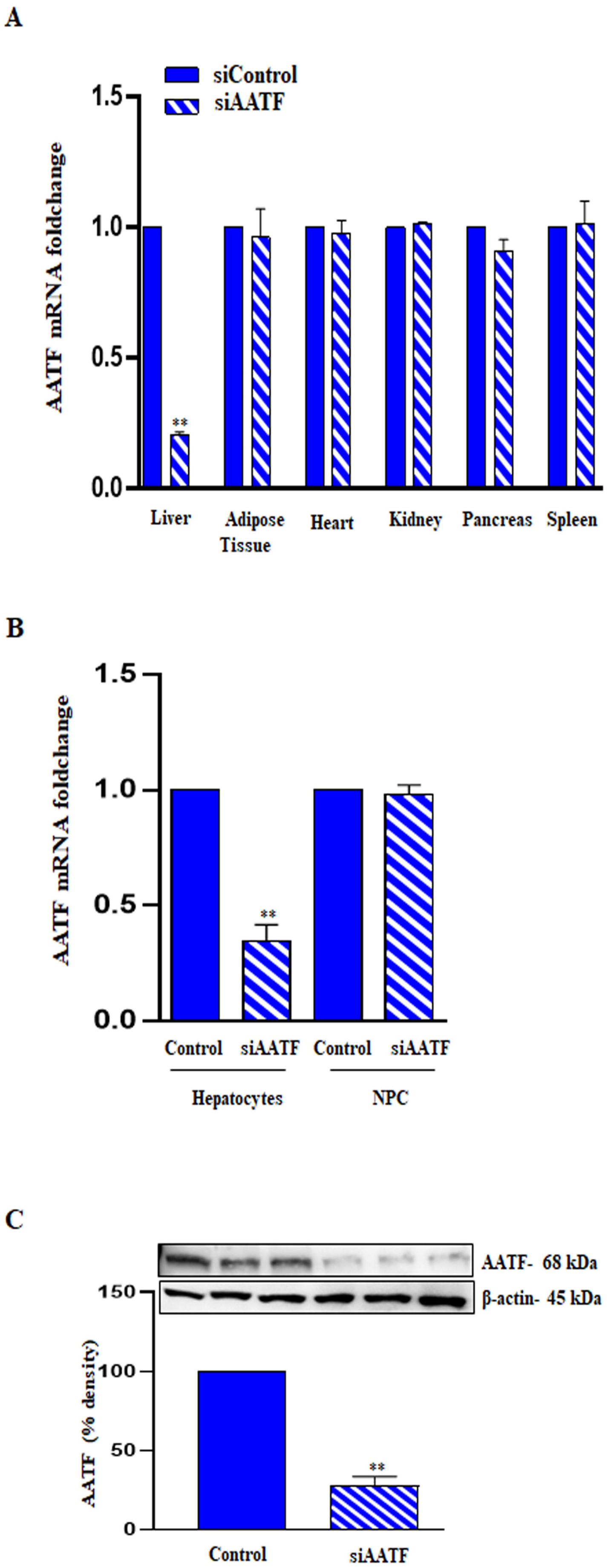
A
siControl
siAATF
AATF mRNA foldchange
1.5
1.0
0.5
0.0
**
Liver
Adipose Tissue
Heart
Kidney
Pancreas
Spleen
B
AATF mRNA foldchange
1.5
1.0
0.5
0.0
**
Control
siAATF
Control
siAATF
Hepatocytes
NPC
C
AATF- 68 kDa
β-actin- 45 kDa
AATF (% density)
150
100
50
0
**
Control
siAATF

**Supplemental Figure 2**

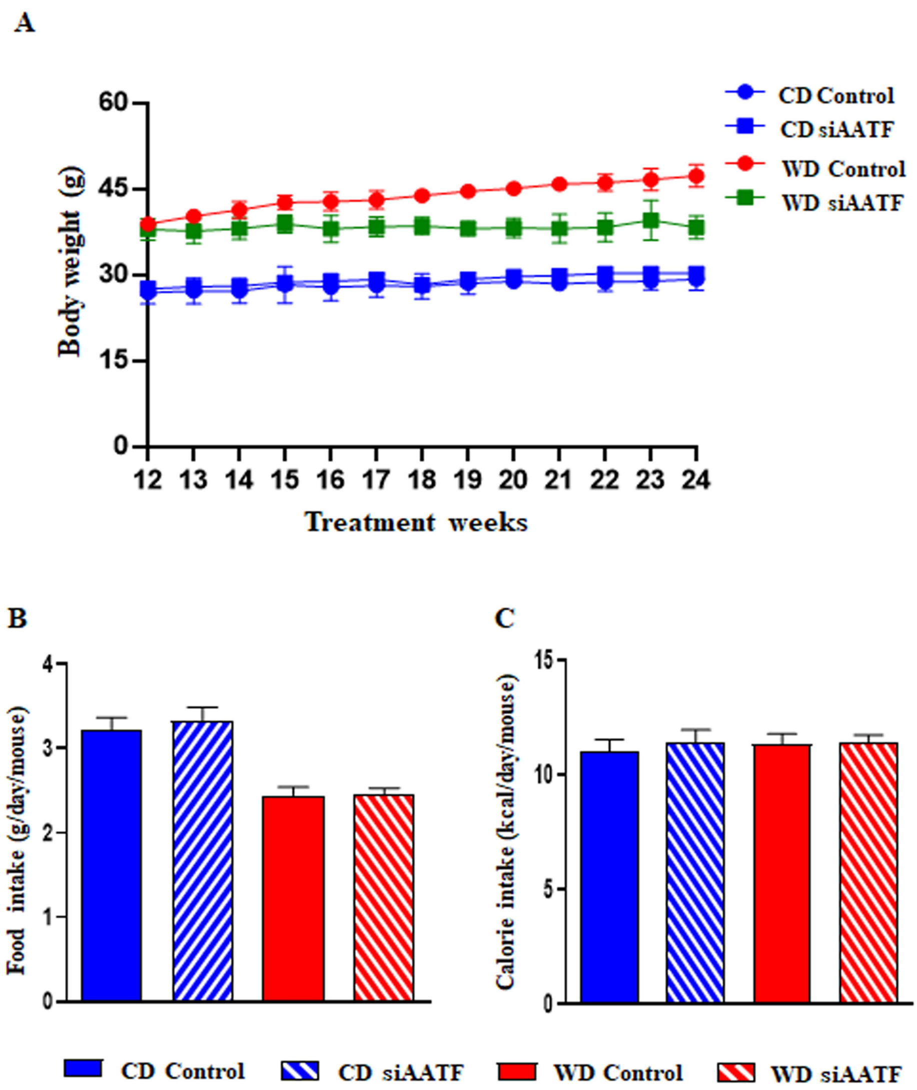

**Supplemental Figure 3**

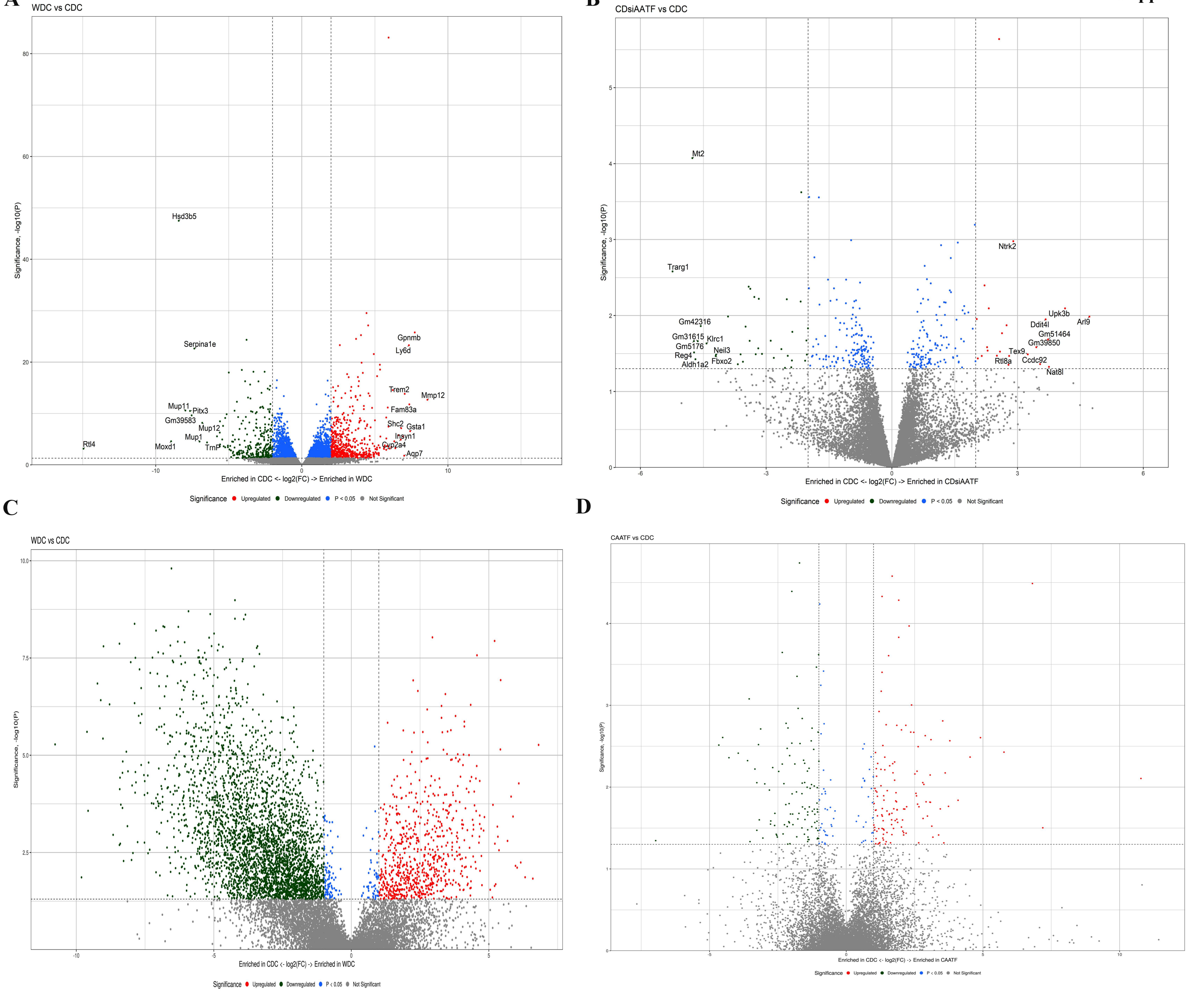

Supplemental Figure 4

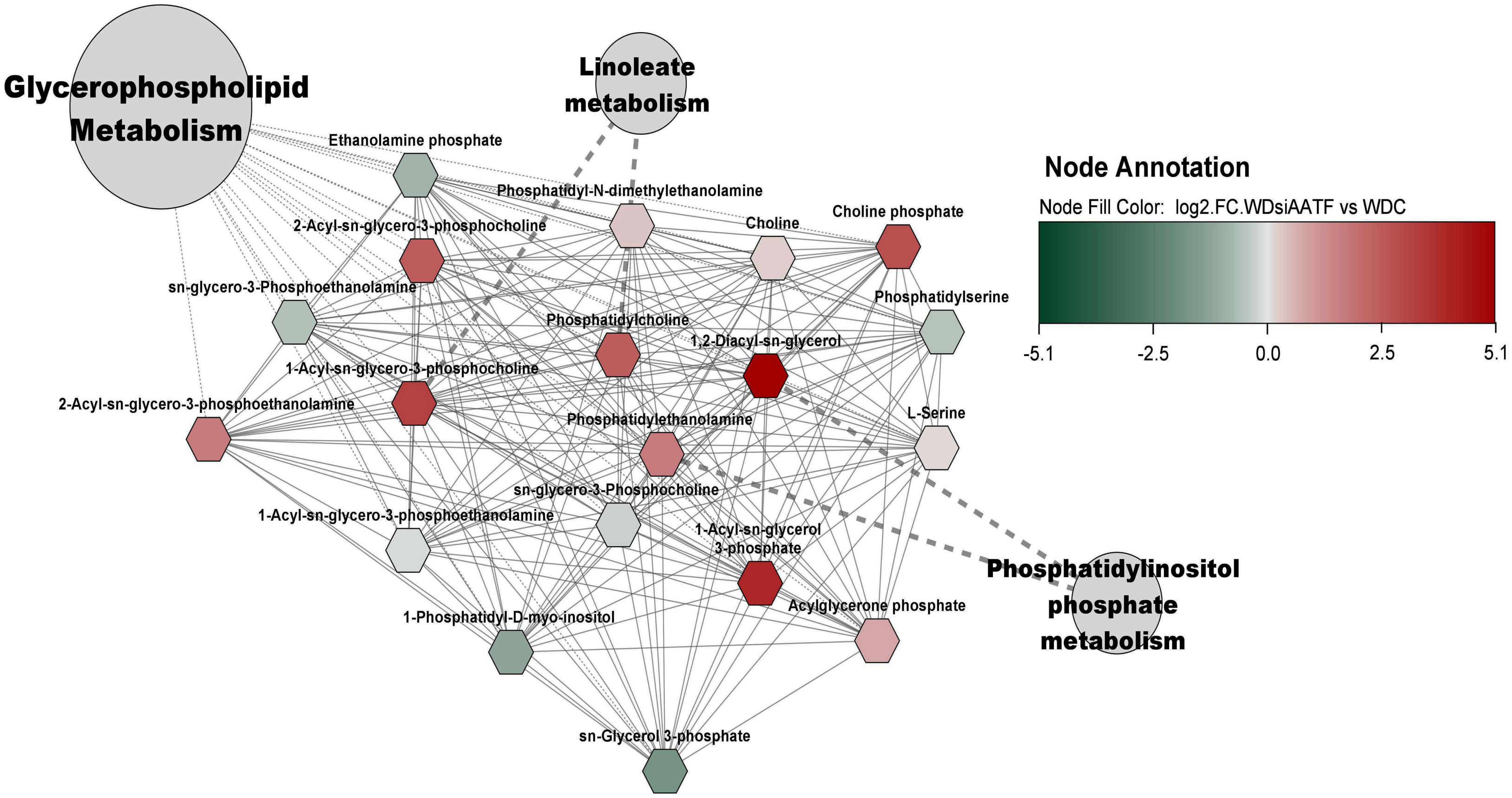

### Supplemental Figure 5

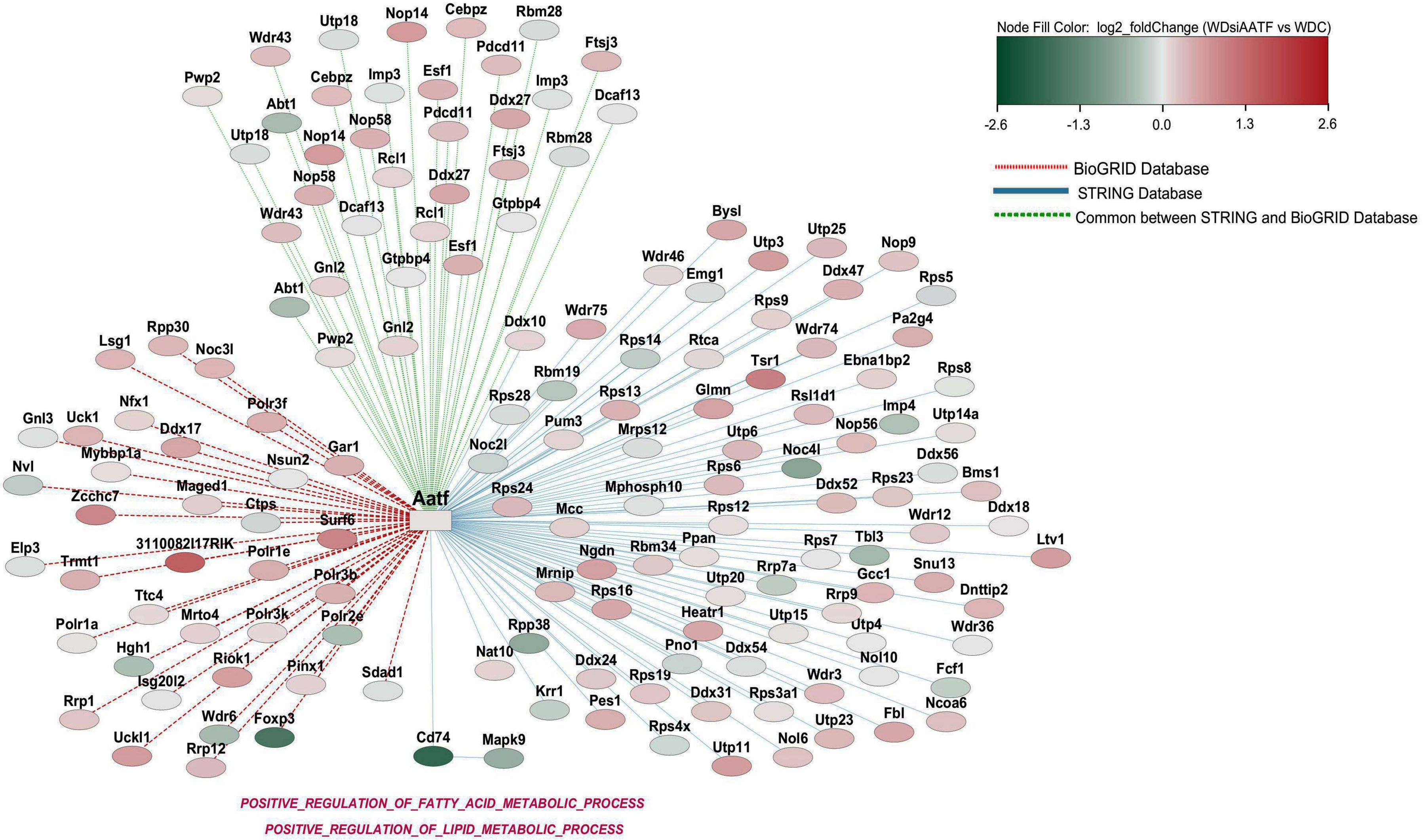

**Supplemental Table 1:** List of primer sequences used in qRT-PCR

| Primer | Sequence |
|---|---|
| **Human AATF** | **F:** GAGTGATGATGCCAGGACGGA<br>**R:** ACTGTCACTTCCCACGGTCTG |
| **Mouse AATF** | **F:** GAGTGATGATGCCAGGACGGA<br>**R:** ACTGTCACTTCCCACGGTCTG |
| **Mouse CHOP** | **F:** GAGCCAGAATAACAGCCGGAAC<br>**R:** ACGTGGACCAGGTTCTGCTTTC |
| **Mouse Grp78** | **F:** TCTTCTCCACGGCTTCCGATA<br>**R:** GTTAGGGGTCGTTCACCTTCAT |
| **Mouse PERK** | **F:** GGAAGAGGACTTTGAGAAAGCA<br>**R:** ACATAGGGCGAGTCTGTCAGTT |
| **Mouse ATF4** | **F:** CCCCTACGACAAGAACATTCA<br>**R:** CACTGGCAGAGTAGTCGAAGG |
| **Mouse IL-1β** | **F:** TGCCACCTTTTGACAGTGATGA<br>**R:** TGATGTGCTGCTGCGAGATTTG |
| **Mouse IL-6** | **F:** GGAGCCCACCAAGAACGATA<br>**R:** AACTGGATGGAAGTCTCTTGC |
| **Mouse TNF-α** | **F:** TAGCCCACGTCGTAGCAAACC<br>**R:** CTTTGAGATCCATGCCGTTGGC |
| **Mouse Col1A1** | **F:** TGACTGGAAGAGCGGAGAGTA<br>**R:** AGACGGCTGAGTAGGGAACA |
| **Mouse Col3A1** | **F:** GACCTAAGGGCGAAGATGGC<br>**R:** GAAGCCACTAGGACCCCTTTC |
| **Mouse α-SMA** | **F:** CTACTGCCGAGCGTGAGATTGT<br>**R:** CCCGCTGACTCCATCCCAATGA |
| **Mouse TGF-β** | **F:** GCTGCATATCGTCCTGTGG<br>**R:** CTTCCATTTCCACATCCGACT |

| | |
|---|---|
| **Mouse FASN** | **F:** TGGGTGGGTGTGAGTGGTTC<br>**R:** GGGCAATGCTTGGTCCTTTGA |
| **Mouse ACC1** | **F:** GGCAGCTCTGGAGGTGTATGT<br>**R:** TGGGATGTGGGCAGCATGAA |
| **Mouse ACC2** | **F:** GTATGGCGTCCCTGGAGGTTTA<br>**R:** CGGTGCTGCAGGCTGTTTAG |
| **Mouse SCD1** | **F:** CCCCTACGGCTCTTTCTGAT<br>**R:** TGGTCACGAGCCCATTCATA |
| **Mouse CPT1α** | **F:** GGCTTCCATGACTCGGCTCTT<br>**R:** ACCTCTGCTCTGCCGTTGTT |
| **Mouse ACOX1** | **F:** TGCAGACGGCCAGGTTCTT<br>**R:** CTGGCTCGGCAGGTCATTCA |
| **Mouse PGC1α** | **F:** GGATGGTGCACAGAACCGATA<br>**R:** CCAACCAGAGCAGCACACTAT |
| **Mouse ACSL1** | **F:** GAAGGCAGCTCCATCTAATCA<br>**R:** TCTCTATGCAGAATTAGTTCCAA |
| **Mouse SREBP1** | **F:** CCAGCAGGTCCCAGTTGTACT<br>**R:** TCACGGTGGCTCCTGCATCT |
| **Mouse LCAD** | **F:** CCAGGAACTACGTGAAGCAAAG<br>**R:** TTATGCTGCACCGTCTGTATGT |
| **Mouse PNPLA2** | **F:** GAAGCTGCTGTGGTGGAGGA<br>**R:** CCACTCCAACAAGCGGATGG |
| **Human β Actin** | **F:** AGAGATGGCCACGGCTGCTT<br>**R:** CAGGACTCCATGCCCAGGAA |
| **Mouse β Actin** | **F:** CAGCCTTCCTTCTTGGGTATGG<br>**R:** CCTGCTTGCTGATCCACATCT |
